# Ubiquitination regulates cytoophidium assembly in *Schizosaccharomyces pombe*

**DOI:** 10.1101/2021.12.06.470909

**Authors:** Christos Andreadis, Tianhao Li, Ji-Long Liu

## Abstract

CTP synthase (CTPS), a metabolic enzyme responsible for the *de novo* synthesis of CTP, can form filamentous structures termed cytoophidia, which are evolutionarily conserved from bacteria to humans. Here we used *Schizosaccharomyces pombe* to study the cytoophidium assembly regulation by ubiquitination. We tested the CTP synthase’s capacity to be epigenetically modified by ubiquitin or be affected by the ubiquitination state of the cell, showed that CTPS is immunoprecipitated with ubiquitin, and that ubiquitination is important for the maintenance of the CTPS filamentous structure in fission yeast. We have identified proteins which are in complex with CTPS, including specific ubiquitination regulators which significantly affect CTPS filamentation, and mapped probable ubiquitination targets on CTPS. Furthermore, we discovered that a cohort of deubiquitinating enzymes is significant for the regulation of cytoophidium morphology. Our study provides a framework for the analysis of the effects that ubiquitination and deubiquitination have on the formation of CTPS filaments.

## Introduction

CTP synthase (CTPS) is the enzyme that catalyzes the rate-limiting step in the production of CTP from UTP in the *de novo* synthesis of CTP (Ozier-Kalogeropoulos et al., 1991, Ozier-Kalogeropoulos et al., 1994). Dysregulation of CTP pool homeostasis and increased CTPS levels have been correlated with a number of cancers, rendering CTPS an important target for drug development (Williams et al., 1978, Whelan et al., 1993, Groblewski et al., 1995, Hatse et al., 1999, Verschuur et al., 2000, Verschuur et al., 2001, Verschuur et al., 2002, Furuta et al., 2010, Wise and Thompson, 2010, Nilsson et al., 2014). Notably, CTPS is suggested to play a functional role in cancer metabolism, since knocking down of the enzyme in Drosophila cancer models is correlated with the reduction of tumour formation (Willoughby et al., 2013).

A characteristic property of CTPS is the assembly into membraneless filamentous structures, termed cytoophidia, which is conserved among evolutionarily divergent species including bacteria (Ingerson-Mahar et al., 2010), archaea (Zhou et al., 2020), budding yeast (Noree et al., 2010), fission yeast (Zhang et al., 2014), *Drosophila* (Liu, 2010), and mammalian cells (Carcamo et al., 2011). The biological function of the filaments remains elusive; however, multiple physiological functions have been suggested (reviewed in (Liu, 2011, Aughey et al., 2016, Liu, 2016)). Importantly, the compartmentation of CTPS potentially contributes to metabolism regulation (Aughey et al., 2014, Barry et al., 2014, Noree et al., 2014, Strochlic et al., 2014), to coping under stress (Aughey et al., 2014), to cell proliferation (Tastan and Liu, 2015, Aughey et al., 2016), along with having cytoskeleton-like functions (Ingerson-Mahar et al., 2010). Even though the progress of CTPS filament research has been significant, the modes of regulation of cytoophidia formation are still relatively obscure (Ingerson-Mahar et al., 2010, Liu, 2010, Carcamo et al., 2011, Chen et al., 2011, Azzam and Liu, 2013, Gou et al., 2014).

In the past few years, a connection has been gradually established between CTPS filamentation and the ubiquitination system. The ubiquitin E3 ligase activity of the proto-oncogene Cbl has been found to be required for CTPS filamentation, without affecting CTPS protein levels in *Drosophila* ovaries during endocycles (Wang et al., 2015). It is not yet clear whether this effect of ubiquitination on cytoophidia formation is direct, or whether there is a still unknown component of the CTPS complex whose regulation by ubiquitination affects the filamentation (Pai et al., 2016). Furthermore, a degenerate cohort of fission yeast membrane trafficking deubiquitinating enzymes (DUBs) which were found to mediate cell polarity and survival also seem to target CTPS (Kouranti et al., 2010, Beckley et al., 2015), thus pointing to a possible regulatory role of deubiquitination in CTPS filamentation.

Previous studies in *S. cerevisiae* and mammalian cells have shown that CTPS can be regulated via phosphorylation by kinases (Park et al., 1999, Choi et al., 2003, Park et al., 2003, Han et al., 2005, Chang et al., 2007). We have recently shown that TOR pathway mediates cytoophidium assembly in mammalian and fission yeast cells (Andreadis et al., 2019, Sun and Liu, 2019b). In addition to phosphorylation, ubiquitination seems to have an involvement in the cytoophidium assembly in *Drosophila* and mammalian cells (Pai et al., 2016). In these cases, CTPS ubiquitination is negatively associated with cytoophidia formation, while inhibition of ubiquitination leads to a reduction of cells containing cytoophidia under glutamine deprivation (Peng et al., 2003, Pai et al., 2016).

Here, we used *S. pombe* to study the capability of ubiquitination to affect the regulation of CTPS cytoophidium formation. By using drugs that modulate ubiquitination and deubiquitination efficiency, as well as by using a series of ubiquitin ligase and DUB deletion mutants, we built upon previous results to show that ubiquitination, as a proteasome-independent modification, significantly affects CTPS cytoophidium formation, while improper deubiquitination significantly affects the morphology of CTPS filaments. Further to this, we performed protein interaction analyses to identify the proteins in complex with CTPS, while also focusing on specific CTPS residues, as potential ubiquitination targets. Our data showcase the importance of ubiquitination in regulating CTPS filamentation, while extending the spectrum of organisms in which CTPS filamentation seems to be modulated by ubiquitination.

## Results

### Ubiquitination and deubiquitination affect CTPS filamentation

We used a *S. pombe* strain that we have previously constructed (Zhang et al., 2014), in which CTPS is fused with YFP tag, to study the effects of ubiquitination in fission yeast cytoophidia. The Cts1–YFP strain has the same growth profile as the WT strain (Zhang et al., 2014). In our model system, cytoophidia are formed in exponential phase, in which 92.7% of the cells on average contain cytoophidia (Figure 1A and 1K), and get degraded in stationary phase, as we have previously shown (Andreadis et al., 2019). We have interfered with the ubiquitination and deubiquitination systems in exponentially growing cells and monitored the changes in CTPS cytoophidium formation. More specifically, we applied E1-ubiquitin ligase inhibitor PYR-41, proteasome inhibitor MG132, and DUB inhibitor PR619 in logarithmic cultures for 2 hours at a range of concentrations, to study the effect on CTPS filamentation (Figure 1B-J). When the ubiquitination system was impaired by PYR-41 E1-ubiquitin ligase inhibitor (Figure 1B-D), we observed a strong and significant reduction of cytoophidia, correlated to the concentration of the inhibitor applied, ranging from 37.9% (50μM PYR-41, p<0.001) to 89.2% (75μM PYR-41, p<0.0001) and 97.4% (100μM PYR-41, p<0.0001), compared to the control strain (Figure 1K).

**Fig 1.**
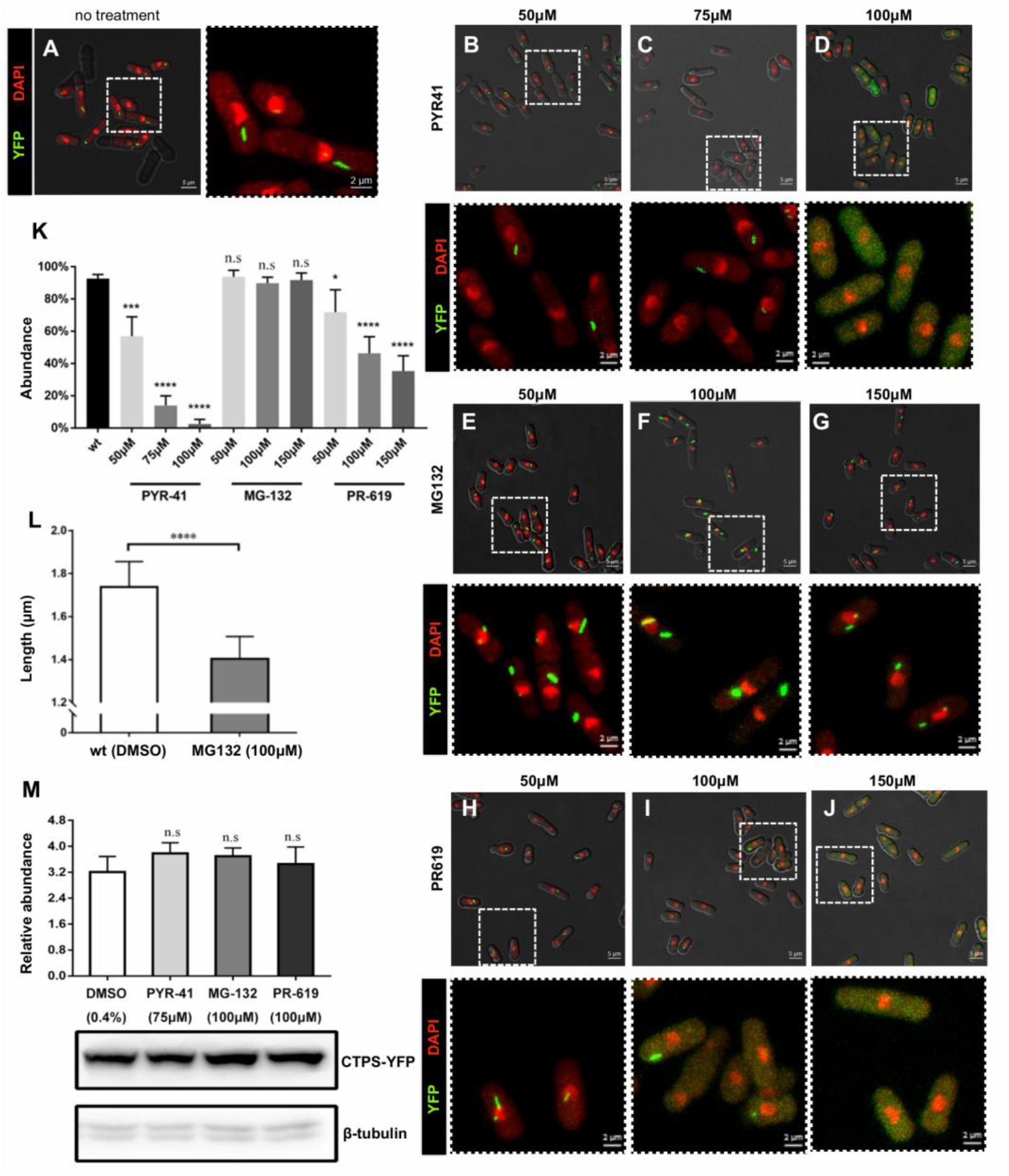
Ubiquitination maintains the filamentous structure of CTP synthase in *S. pombe*. (A) CTP synthase fused to YFP was expressed endogenously under the control of *Cts1* native promoter. *S. pombe* control cells were cultured until exponential phase (OD600=0.4) in YE4S rich medium with the addition of 0.4% DMSO for 2h, fixed, and observed by confocal microscopy. An overlap image of brightfield (white), YFP (green), and DAPI (red) is presented on the left part (scale bar=5μm), while a magnified image of the dotted rectangle is presented on the right with an overlap of the YFP and DAPI channels for clarity (scale bar=2μm). (B-D) Exponentially growing *S. pombe* cells (OD600=0.4) were treated with 50μM, 75μM, 100μM PYR-41 for 2h, fixed, and observed by confocal microscopy as in panel A. (E-G) Exponentially growing *S. pombe* cells (OD600=0.4) were treated with 50μM, 100μM, 150μM MG-132 for 2h, fixed, and observed by confocal microscopy as in panel A. (H-J) Exponentially growing *S. pombe* cells (OD600=0.4) were treated with 50μM, 100μM, 150μM PR-619 for 2h, fixed, and observed by confocal microscopy as in panel A. (K) Quantification of the cells with visible cytoophidia is plotted for the conditions described in panels A-J expressed as percentage of cells containing cytoplasmic cytoophidia in each of the conditions tested. Error bars show the mean ± S.D.: as calculated from three independent experiments (>400 cells were manually counted per strain per trial, ****: *p*<0.0001, ***: *p*<0.001, *: *p*<0.05, n.s.: not significant). (L) Exponentially growing *S. pombe* cells (OD600=0.4) were treated with 0.4% DMSO or 100μM MG-132 for 2h, and the average length of cytoplasmic cytoophidia was calculated. Error bars show the mean ± S.D.: as calculated from three independent experiments (>400 cells were manually counted per strain per trial, ****: *p*<0.0001). (M) Protein extracts of exponentially growing cultures of control cells (DMSO) and of cells grown upon PYR-41 (75μM), MG-132 (100μM), and PR-619 (100μM) for two hours, were analysed by Western blot. The quantification of the Cts1-YFP protein is presented in the upper panel as relative Cts1 protein abundance, after normalization over the levels of β-tubulin. Error bars show the mean ± S.D.: as calculated from three independent experiments (n.s.: *p*>0.05). The bottom panel shows a representative image of the Western blot membrane, along with the β-tubulin levels.

The percentage of cells containing cytoophidia was not affected by the impairment of proteasome, since the proteasome inhibitor MG-132 did not result to any significant change in the cytoophidium formation, ranging from 4.6% increase (50μM MG-132, p>0.5) to 3.2% decrease (100μM MG-132, p>0.5) and 2.8% decrease (150μM MG-132, p>0.5), compared to the control strain (Figures 1E-G, 1K). However, the length of cytoophidia was significantly reduced under this condition from 1.73μM to 1.41μM on average (p<0.0001) (Figure 1L).

When the deubiquitination system was compromised by PR-619 DUB inhibitor (Figures 1H-J), the number of cells containing cytoophidia was significantly reduced in a drug dose-dependent manner, ranging from 14.3% (50μM PR-619, p<0.05) to 32% (100μM PR-619, p<0.0001) and 61.7% (150μM PR-619, p<0.0001), compared to the control strain (Figure 1K). Interestingly, prolonged treatment of the cells with E1-ubiquitin ligase inhibitor PYR-41, proteasome inhibitor MG132, and DUB inhibitor Pr619 for 4.5 hours, showed a sufficient recovery of the phenotype (Figure S1). Assuming that the activity of the drugs remains unchanged for the duration of the prolonged treatment, this result highlights the dynamic nature of CTPS cytoophidia, as well as their capability to adapt under this type of stress.

After observing the substantial dependency of CTPS filamentation on the ubiquitination and deubiquitination systems in fission yeast, we sought to investigate whether this effect could be attributed to a possible change in the CTPS protein levels. Our expression analyses showed that this was not the case, as the CTPS protein levels did not change significantly upon impairment of ubiquitination and deubiquitination systems or upon inhibition of the proteasome pathway (p>0.15) (Figure 1M).

Taken together, our data show that the ubiquitination state of the cell plays an important role in maintaining the filamentous structures of CTPS in fission yeast, which are naturally formed under physiological growth conditions.

### *S. pombe* CTPS immunoprecipitates with ubiquitin

To explore whether the effect of ubiquitination on cytoophidium formation is direct, we conducted co-immunoprecipitation (co-IP) experiments to determine whether CTPS can be ubiquitinated. We constructed a fission yeast strain for the co-IP analyses in which CTPS is tagged with the small tags HA and flag. The strain Cts1-flag-3HA strain provided us with the flexibility of conducting IP experiments using small tags, which are non-eukaryotic and do not interfere with the complex assembly or the protein function. Our co-IP experiments showed that CTPS immunoprecipitates with ubiquitin (Figure 2A), and a shift in the Cts1 protein size, corresponding to the small size of ubiquitin (<10kDa) (Figure 2B). This finding is indicative of a potential direct effect of ubiquitination on CTPS compartmentation, though other proteins in complex with CTPS may also be ubiquitinated.

**Fig 2.**
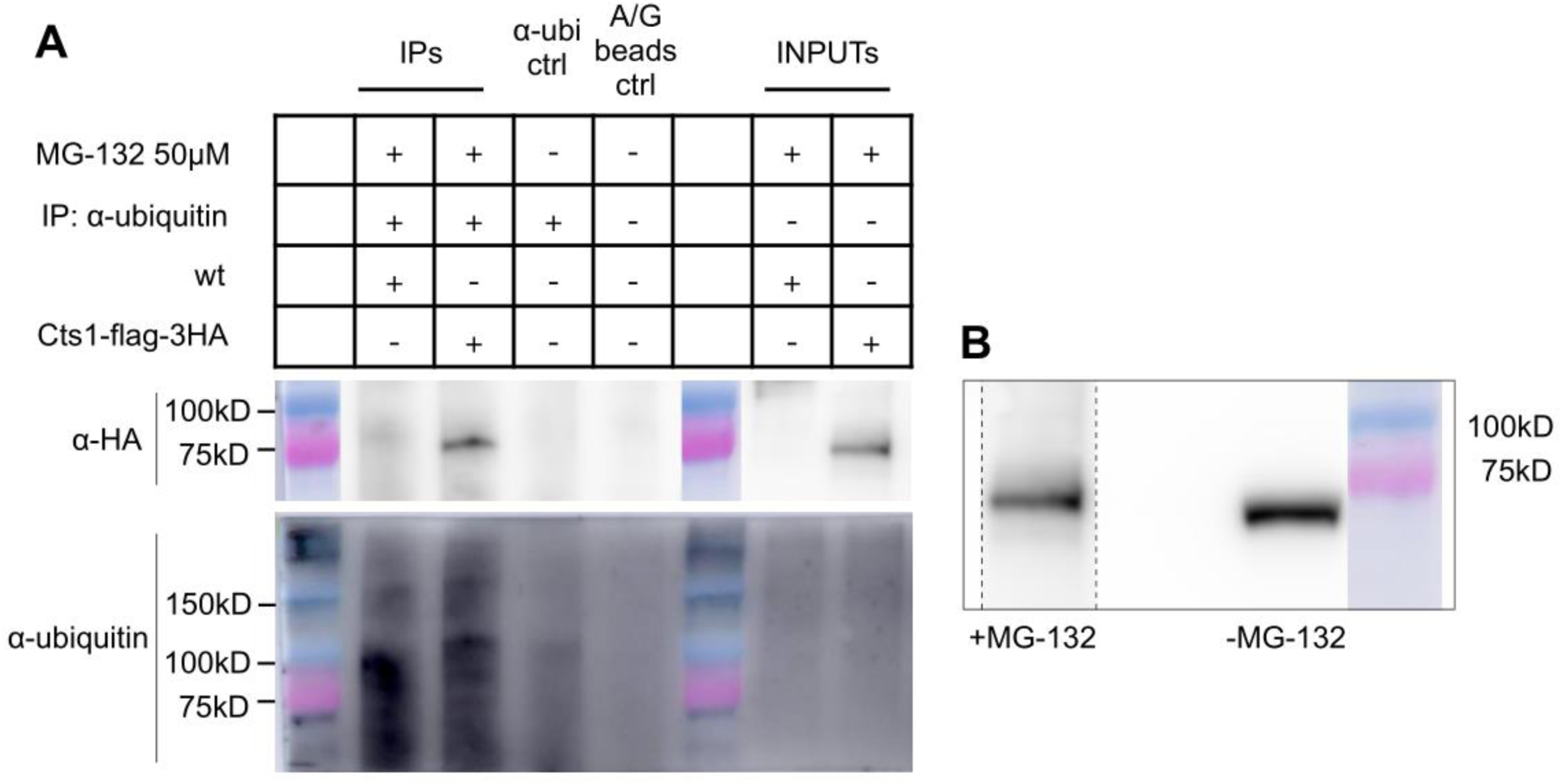
Cts1 immunoprecipitates with ubiquitin. (A) Cts1-flag-3HA and wild type (wt) cells were cultured in rich YE4S medium until they reached exponential phase (OD_600_=0.4), followed by protein extraction. The protein extract was subsequently immunoprecipitated using α-ubiquitin antibody, followed by incubation with A/G beads. The samples were then electrophoresed on SDS-PAGE and the membrane probed firstly by α-ΗΑ antibody and, after stripping, by α-ubiquitin antibody. Information about the protein extract ran on each lane is given in the upper part of the figure for immunoprecipitated samples (IPs) and input samples (INPUTs). Control samples included one for the α-ubiquitin antibody background (α-ubi ctrl), one for the A/G beads background (A/G beads ctrl), in addition to the wild type (wt) strain (no tag). MG132 proteasome inhibitor was used to ensure no protein degradation. (B) Western blot analysis of INPUT sample of Cts1-flag-3HA strain when proteasome is inhibited (+MG-132) (left), as in (A), probed by α-HA antibody. Note the shift observed to the band of Cts1-flag-3HA (left) (dotted part of membrane exposed for 2 mins), compared to a TCA-extracted protein sample obtained under normal growth conditions (- MG132) (right) (same membrane exposed for 3 seconds).

### CTPS cytoophidium formation is affected by ubiquitin ligases

We used the Cts1-flag-3HA strain and performed a series of FLAG-HA tandem affinity purification assays (TAP assays), combined with analyses by liquid chromatography and mass spectrometry (LC-MS), in order to characterise the fission yeast CTPS protein complex. We chose to use the HA-FLAG TAP assay because it involves a double sequential elution which takes advantage of the double tag on the protein, offering higher specificity and pull-down of fewer contaminants, compared to other systems. This strategy offers a very low rate of cross-reactivity with non-targeted endogenous proteins, while the tags do not sterically affect the bait proteins’ interactions. Our analyses showed a number of proteins that are in complex with CTPS. Based on the biological processes in which they participate, these proteins are either structural, or involved in metabolism, transport or stress (Table S1). Further to this, we used Cytoscape software (Cline et al., 2007) to visualise the potential networks in which the proteins in complex with CTPS participate, along with the biological networks gene ontology tool (BiNGO) (Maere et al., 2005) to determine the gene ontology (GO) terms that are significantly overrepresented in our group of proteins (Figure S2). This analysis, not only mapped *S. pombe* CTPS interactome, but also provided us with a mechanistic insight into the regulation of cytoophidia formation by ubiquitination, by the identification of Mug30 and Ubr11 ubiquitin ligases in complex with CTPS.

We constructed deletion mutant strains for the Mug30 and Ubr11 ubiquitin ligases in a Cts1-YFP background in order to monitor the effects of the lack of these enzymes in the filamentation of CTPS. We performed a detailed confocal microscopic analysis (Figure 3A-C) and found that the abundance of cytoplasmic cytoophidia in *mug30*Δ mutant strains was significantly reduced by 21% (p<0.0001) compared to the Cts1-YFP reference strain (Figure 3D). The average length of CTPS cytoophidia was significantly reduced in both mutants, compared to the Cts1-YFP strain. More specifically the average length was reduced by 21.3% in *mug30*Δ (p<0.01), and by 22.8% in *ubr11*Δ strains (p<0.01) (Figure 3E). We then grouped the cytoophidia of each strain into three groups based on their length (<1μm, between 1-2μm, >2μm) and conducted a more in-depth analysis on how their length changed. We found a strong shift from longer to shorter cytoophidia in all mutants, compared to the control strain (Figure 3F). In particular, the small cytoophidia (<1μm) increased 11.7 folds in *mug30*Δ (p<0.01) and 9.5 fold in *ubr11*Δ strains (p<0.05), as compared to the Cts1-YFP reference strain. In agreement with this trend, the large cytoophidia (>2μm) decreased 2.3-fold in *mug30*Δ (p<0.05), and 5.2-fold in *ubr11*Δ strains (p<0.05), as compared to the Cts1-YFP reference strain (Figure 3F). We then examined whether these trends were related to any changes in the protein levels of CTPS in these mutants and found that there was no significant change (Figure 3G).

**Fig 3.**
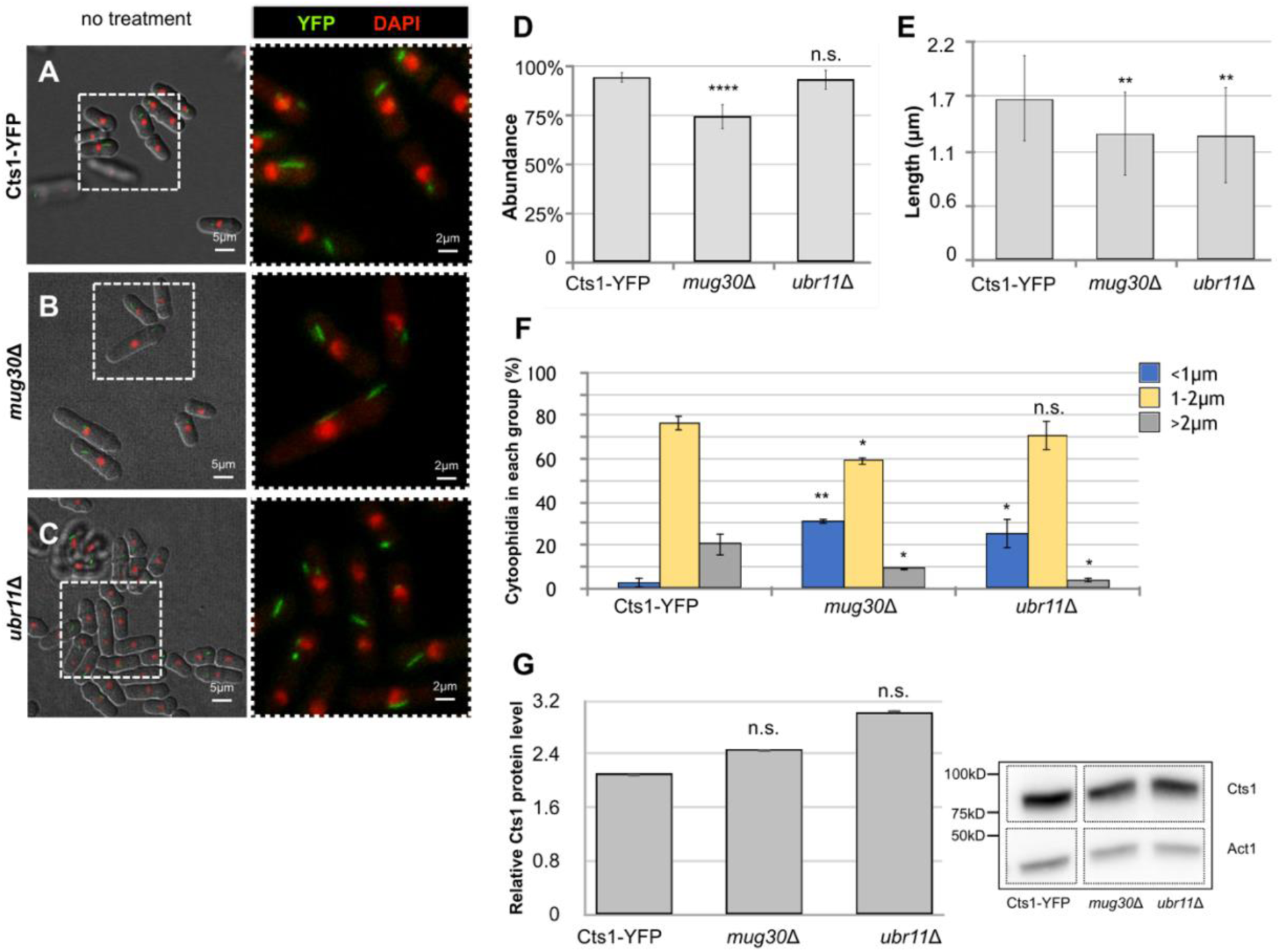
Ubiquitination ligases affect CTPS cytoophidium formation in *S. pombe*. (A-C) CTP synthase fused to YFP was expressed endogenously under the control of *Cts1* native promoter. *S. pombe* control cells (A), as well *mug30*Δ (B), and *ubr1* Δ (C) in a Cts1-YFP background were cultured until exponential phase (OD600=0.4) in YE4S rich medium, fixed, and the presence of cytoophidia was studied by confocal microscopy. An overlap image of brightfield (white), YFP (green), and DAPI (red) is presented on the left part of each image (scale bar=5μm), while a magnified image of the dotted rectangle is presented on the right with an overlap of the YFP and DAPI channels for clarity (scale bar=2μm). (D) Quantification of the cells with visible cytoophidia is plotted for the control and mutant strains described in panels A-C, expressed as percentage of cells containing cytoplasmic cytoophidia in each of the strains tested. Error bars show the mean ± S.D.: as calculated from three independent experiments (>400 cells were manually counted per strain per trial, ****: *p*<0.0001, n.s.: not significant). (E) The average length of cytoplasmic cytoophidia in the mutant cells was calculated and plotted along with the average length of cytoplasmic cytoophidia in control Cts1-YFP cells grown in YE4S rich medium, as in panels A-C. Error bars show the mean ± S.D.: as calculated from three independent experiments (>400 cells were manually counted per strain per trial, **: *p*<0.01). (F) The average length of cytoplasmic cytoophidia as measured in panel E, was used as source to group the filaments into three categories based on their length, as shown. The distribution of cytoophidia, expressed as average percentage of cytoophidia in each category, is plotted for the mutants and the Cts1–YFP control strain. Different color coding is used to distinguish between the three length categories as shown (smaller than 1μm in blue, between 1-2μm in yellow, larger than 2μm in grey). Error bars show the mean ± S.D.: as calculated from three independent experiments (>400 cells were manually counted per strain per trial, **: *p*<0.01, *: *p*<0.05, n.s.: not significant). (G) Cts1-YFP and deletion mutant cells were cultured until exponential phase (OD600=0.4) in YE4S rich medium. Proteins were extracted from equal number of cells, and samples were analyzed by SDS-PAGE Western blot analysis. The Cts1–YFP protein levels were calculated and plotted (left) after normalization over actin levels. A representative image of the Western blot membrane is shown on the right, along with the actin levels. Error bars show the mean ± S.D.: as calculated from three independent experiments (n.s.: not significant). Dotted lines indicate the areas of the membrane that have been cut out.

Collectively, our data introduce some of the protein players involved in the regulation of *S. pombe* CTPS compartmentation by ubiquitination, by identifying specific ubiquitin ligases in complex with CTPS, the action of which significantly affects the abundance and the length of CTPS cytoophidia.

### CTPS filamentation is disrupted in K430R and K264R mutants

Next, we screened *S. pombe* CTPS amino acids based on their ability to become ubiquitinated. We have initially utilised UbiPred software to predict which lysine residues are more probable to be modified by ubiquitin (Radivojac et al., 2010). We have then modelled the three-dimensional conformation of *S. pombe* CTPS, based on the already solved CTPS structure in mammalian cells, by using Phyre2 software (Kelley et al., 2015). After combining the results of the two analyses, we have identified two lysine residues, namely K430 and K264, which, based on their position in the three-dimensional reconstruction of CTPS, are freer to interact with other proteins and have higher probability to become ubiquitinated (Figure 4A).

**Fig 4.**
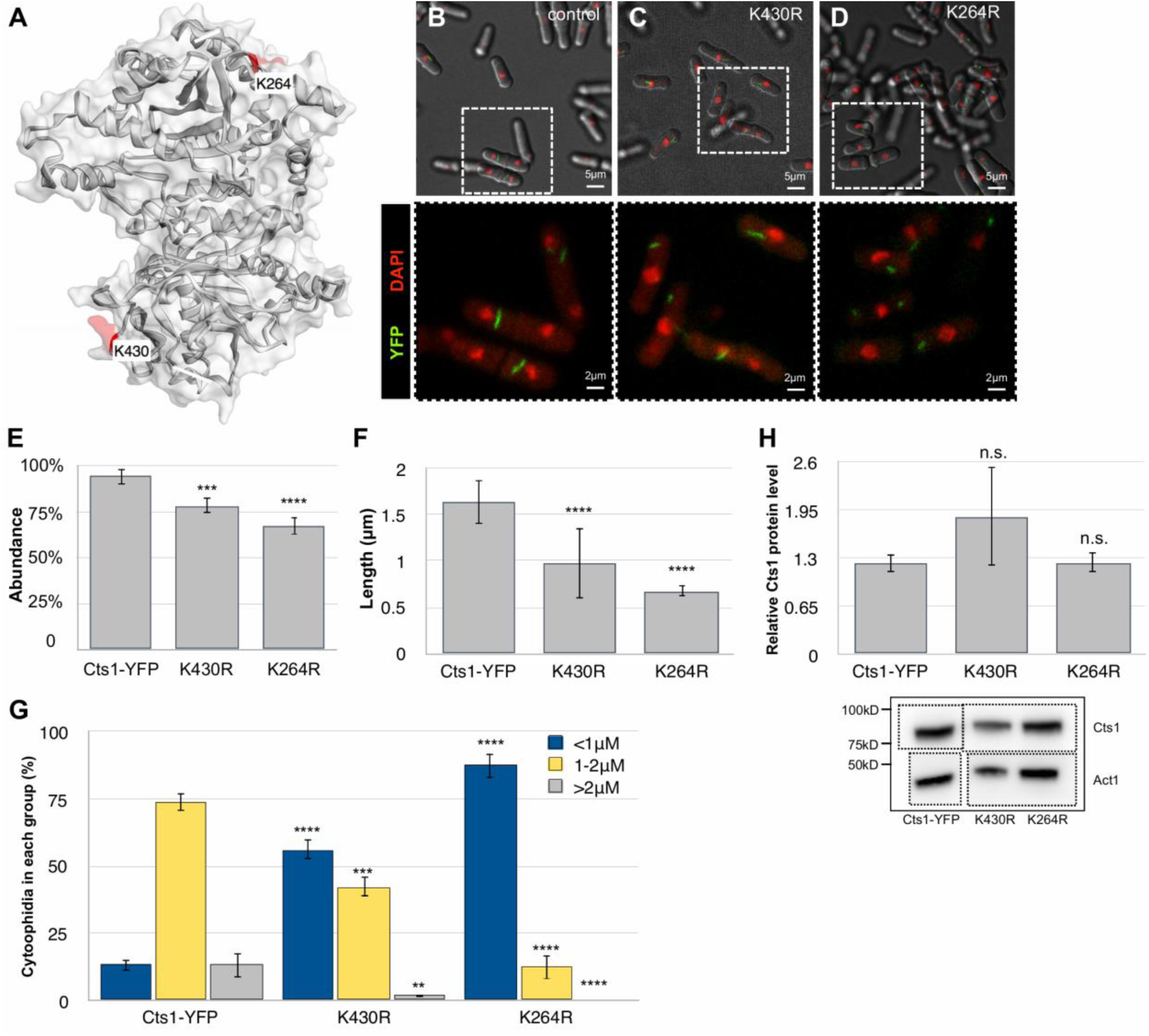
K430R and K264R disrupt CTPS filamentation in *S.pombe*. (Α) Three-dimensional conformation of *S. pombe* CTPS obtained by using Phyre2 and EzMol software, based on the solved conformation of mammalian CTPS. *In silico* analysis by UbiPred software showed that K430 and K264 residues are potential ubiquitination targets; the positioning of the two amino acids is indicated, along with the surfaces occupied by the residues (in red). (B-D) Confocal microscopy of control (Cts1-YFP) (B), K430R (C) and K264R (D) strains in a Cts1-YFP background, after being cultured until exponential phase (OD600=0.4) in YE4S rich medium and fixed. An overlap image of brightfield (white), YFP (green), and DAPI (red) is presented on the top part of each image (scale bar=5μm), while a magnified image of the dotted rectangle is presented at the bottom part, with an overlap of the YFP and DAPI channels for clarity (scale bar=2μm). (E) Quantification of the cells with visible cytoophidia is plotted for the control, K430R, and K264R strains, expressed as percentage of cells containing cytoplasmic cytoophidia in each of the strains tested. Error bars show the mean ± S.D.: as calculated from three independent experiments (>400 cells were manually counted per strain per trial, ****: *p*<0.0001, ***: *p*<0.001). (F) The average length of cytoplasmic cytoophidia in K430R and K264R mutant cells was calculated and plotted along with the average length of cytoplasmic cytoophidia in control Cts1-YFP cells grown in YE4S rich medium, as in panels B-D. Error bars show the mean ± S.D.: as calculated from three independent experiments (>400 cells were manually counted per strain per trial, ****: *p*<0.0001). (G) The average length of cytoplasmic cytoophidia as measured in panel F, was used as a source to group the filaments into three categories based on their length, as shown. The distribution of cytoophidia, expressed as average percentage of cytoophidia in each category, is plotted for the mutants and the Cts1–YFP control strain. Different color coding is used to distinguish between the three length categories as shown (smaller than 1μm in blue, between 1-2μm in yellow, larger than 2μm in grey). Error bars show the mean ± S.D.: as calculated from three independent experiments (>400 cells were manually counted per strain per trial, ****: *p*<0.0001, ***: *p*<0.001 **: *p*<0.01). (H) Cts1-YFP, K430R, and K264R mutant cells were cultured until exponential phase (OD600=0.4) in YE4S rich medium. Proteins were extracted from equal number of cells, and samples were analyzed by SDS-PAGE Western blot analysis. The Cts1–YFP protein levels were calculated and plotted (top) after normalization over actin levels. A representative image of the Western blot membrane is shown at the bottom, along with the actin levels. Error bars show the mean ± S.D.: as calculated from three independent experiments (n.s.: not significant). Dotted lines indicate the areas of the membrane that have been cut out.

Following this, we proceeded with mutating each of these lysine residues into arginine, rendering them incapable of being ubiquitinated. Subsequently, we constructed the fission yeast Cts1-YFP strains bearing either K430R or K264R point mutations on CTPS and studied how this could affect the regulation of cytoophidia formation by performing confocal microscopy (Figure 4B-D). We found that, compared to the reference Cts1-YFP strain, the abundance of cytoophidia was significantly reduced in both K430R and K264R mutants, by 16.7% in (p<0.001), and by 28.1% in (p<0.0001), respectively (Figure 4E).

The effect of the mutations on cytoophidia formation was more evident on their average length, which was reduced by 40.3% in K430R cells (p<0.0001) and by 58.3% in K264R cells (p<0.0001), compared to the Cts1-YFP reference strain (Figure 4F). Furthermore, when we grouped the CTPS cytoophidia into three groups based on their length (<1μm, between 1-2μm, >2μm), our results showed a significant 4.3-fold increase of smaller cytoophidia (<1μm) in K430R strain (p<0.0001) and an equally significant 6.7-fold increase of smaller cytoophidia in K264R strain (p<0.0001) as compared to the Cts1-YFP strain (Figure 4G). In agreement with the trend of the reduction of cytoophidia length in the point mutation strains, we observed an 8.2-fold reduction in large cytoophidia (>2μm) in K430R cells, (p<0.01), and a 13.2-fold reduction in K264R cells (p<0.0001), as compared to the reference Cts1-YFP strain (Figure 4G). The phenotypes exhibited by the CTPS cytoophidia in the K430R and K264R mutant strains was not due a change in the protein levels of CTPS, since these were found to be non-significant (Figure 4H).

We have found that point mutations such as K430R and ubiquitin ligase deletion mutants *mug30*Δ and *ubr11*Δ, significantly reduce the strength by which CTPS immunoprecipitates with ubiquitin (6.2-fold, 3.4-fold, 7.4-fold, respectively) (p<0.01) (Figure S3).

Taken together, our results strongly indicate that not only the ubiquitination state of the cell affects cytoophidia formation, but there are specific ubiquitin ligases in complex with CTPS, as well as particular CTPS residues which are involved in the regulation of CTPS compartmentation. Our *in silico* analyses further showed that, apart from K430 and K264 residues, there are additional candidates which could be potential ubiquitination targets, albeit with a lower degree of confidence, nevertheless worth-exploring in a future study (Figure S4).

### Deubiquitinating enzymes affect CTPS cytoophidia morphology

Previous studies focused on cell polarity and survival have incidentally demonstrated, by employing comparative proteomics, biochemistry and microscopy, that a series of five degenerate deubiquitinating enzymes target CTPS (Beckley et al., 2015). These genes encode four ubiquitin C-terminal hydrolases (Ubp4, Ubp5, Ubp9, Ubp15) and one ubiquitin-specific protease (Sst2). We wanted to examine whether the deubiquitinating function of these proteins is necessary for the CTPS compartmentation and whether it affects in any way the CTPS filament formation.

To address this question, we constructed a series of single and multiple mutants of the deubiquitinating genes and studied the effects on CTPS filamentation by confocal microscopy (Figure 5A-H). We did not observe any significant change of the percentage of cells containing cytoophidia in *sst2*Δ, *ubp4*Δ, *ubp5*Δ, *ubp9*Δ, *ubp15*Δ, *sst2*Δ *ubp4*Δ *ubp9*Δ *ubp15*Δ, and *sst2*Δ *ubp4*Δ *ubp5*Δ *ubp9*Δ mutants (Figures 5B-H), as compared to the control strain (Figure 5A). The changes in the abundance of cytoophidia were insignificant (Figure 5I). However, the average length of cytoplasmic cytoophidia in the series of single and multiple deubiquitinating mutants was significantly reduced. More specifically, while the average length of cytoplasmic cytoophidia in the WT (Cts1-YFP) was 1.77μm (±0.13μm), our calculations showed a significant change in the average cytoophidia length which was 1.46μm (±0.03μm) in *sst2*Δ (p<0.0001), 1.56μm (±0.13μm) in *ubp4*Δ (p<0.001), 1.46μm (±0.05μm) in *ubp5*Δ (p<0.0001), 1.51μm (±0.08μm) in *ubp9*Δ (p<0.0001), 1.39μm (±0.06μm) in *ubp15*Δ (p<0.0001), 1.63μm (±0.12μm) in *sst2*Δ *ubp4*Δ *ubp9*Δ *ubp15*Δ (p<0.05), and 1.55μm (±0.19μm) in *sst2*Δ *ubp4*Δ *ubp5*Δ *ubp9*Δ mutant strain (p<0.001) (Figure 5J).

**Fig 5.**
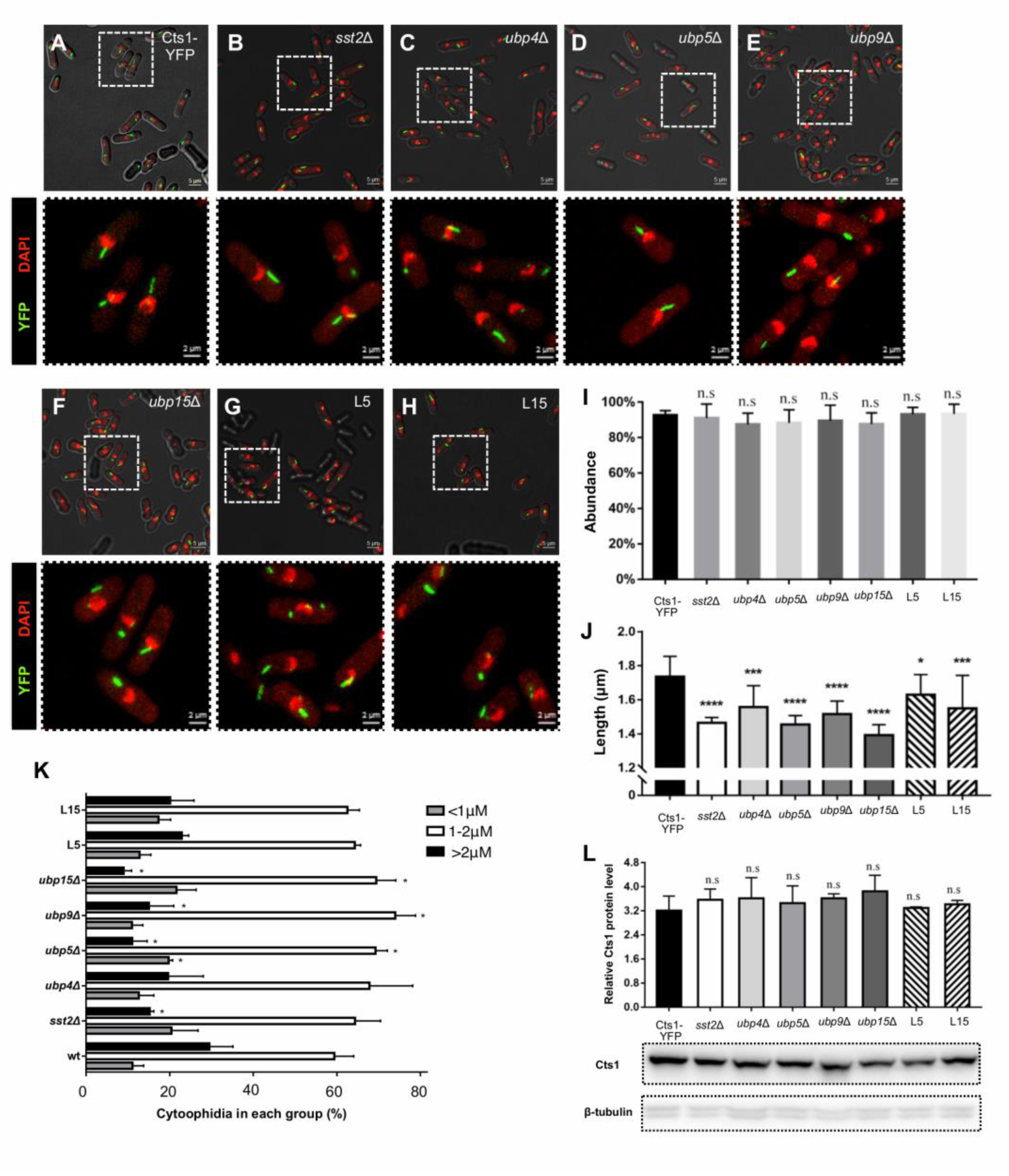
Deubiquitinating mutants affect the morphology of cytoophidia. (A-H) *S. pombe* Cts1-YFP cells were cultured until exponential phase (OD600=0.4) in YE4S rich medium, and fixed. Representative images of confocal microscopic analyses are shown for wild type Cts1-YFP strain (A) and a series of deubiquitinating mutants constructed in a Cts1-YFP background, as shown (B-H). An overlap image of brightfield (white), YFP (green), and DAPI (red) is presented on the top part of each image (scale bar=5μm), while a magnified image of the dotted rectangle is presented at the bottom part, with an overlap of the YFP and DAPI channels for clarity (scale bar=2μm). (I) Quantification of the cells with visible cytoophidia is plotted for the strains noted in A-H, expressed as percentage of cells containing cytoplasmic cytoophidia in each of the strains tested. Error bars show the mean ± S.D.: as calculated from three independent experiments (>400 cells were manually counted per strain per trial, n.s.: not significant). (J) The average length of cytoplasmic cytoophidia in deubiquitinating mutant cells was calculated and plotted along with the average length of cytoplasmic cytoophidia in control Cts1-YFP cells grown in YE4S rich medium, as in panels A-H. Error bars show the mean ± S.D.: as calculated from three independent experiments (>400 cells were manually counted per strain per trial, ****: p<0.0001, ***: p<0.001 *: p<0.05). (K) The average length of cytoplasmic cytoophidia as measured in panel J, was used as a source to group the filaments into three categories based on their length, as shown. The distribution of cytoophidia, expressed as average percentage of cytoophidia in each category, is plotted for the mutants and the Cts1–YFP control strain. Error bars show the mean ± S.D.: as calculated from three independent experiments (>400 cells were manually counted per strain per trial, *: p<0.05). (L) Cts1-YFP and deubiquitinating mutant cells were cultured until exponential phase (OD600=0.4) in YE4S rich medium. Proteins were extracted from equal number of cells, and samples were analyzed by SDS-PAGE Western blot analysis. The Cts1–YFP protein levels were calculated and plotted (top) after normalization over β-tubulin levels. A representative image of the Western blot membrane is shown at the bottom, along with the β-tubulin levels. Error bars show the mean ± S.D.: as calculated from three independent experiments (n.s.: not significant). Dotted lines indicate the areas of the membrane that have been cut out. L5: *sst2*Δ *ubp4*Δ *ubp9*Δ *ubp15*Δ, L15: *sst2*Δ *ubp4*Δ *ubp5*Δ *ubp9*Δ mutant. All strains were constructed in a Cts1-YFP background.

We further examined this phenotype by grouping the cytoophidia in three categories based on their length (<1μm, between 1-2μm, >2μm), and subsequently checking how they are distributed in these categories in all mutant and control strains (Figure 5K). A trend of increasing percentage of shorter (<1μm) cytoophidia in *ubp5*Δ (p<0.05) and decreasing percentage of longer (>2μm) cytoophidia in *sst2*Δ, *ubp5*Δ, *ubp9*Δ, and *ubp15*Δ (p<0.05) was observed, as compared to the wild type strain (Figure 5K). Notably, this effect was not related to any changes in the CTPS protein levels. Our expression analyses showed no significant change of the CTPS protein levels in any of the DUB mutant strains (Figure 5L).

Our results demonstrate the importance of the deubiquitination process in retaining the proper formation of CTPS cytoophidia. Our overall data suggest that ubiquitination is a key post-translational modification affecting the process of filamentation of CTPS, be it directly or indirectly, while its reversibility further contributes to the fine-tuning of the CTPS cytoophidium formation in the cells.

## Discussion

Our study has demonstrated a link between CTPS cytoophidium formation and the ubiquitination/deubiquitination processes. We showed that proper, uninhibited ubiquitination and deubiquitination are key to the assembly and maintenance of CTPS filaments. Inhibition of either process lead to a disruption of the physiological formation of CTPS cytoophidia in *S. pombe* cells. We have constructed a putative map of the proteins in complex with fission yeast CTPS, which included particular ubiquitin ligases. By *in silico* analyses we identified specific lysine residues, potential ubiquitination targets, which we found to be important for the proper compartmentation of CTPS. Furthermore, we showed that the absence of specific deubiquitinating enzymes affects the morphology of cytoophidia by reducing their average length. Our results demonstrate that the system of ubiquitination/deubiquitination could be a conserved means of regulation of cytoophidium filamentation.

Previous studies have shown that ubiquitination regulates CTPS activity by promoting CTPS filament formation in Drosophila ovaries during endocycles (Wang et al., 2015). We have previously shown that in mammalian cells the CTPS compartmentation is negatively associated with CTPS ubiquitination, since cytoophidia formation blocked the CTPS ubiquitination and prolonged its half-life (Sun and Liu, 2019a). However, it has also been shown that ubiquitination is required for CTPS filament formation in Drosophila and mammalian cells, since cytoophidia were completely abolished or strongly reduced, respectively, when E1 ubiquitin ligase was inhibited by PYR41 drug (Pai et al., 2016). These opposing roles that ubiquitination seems to have on cytoophidia formation could be the result of differential effects of ubiquitin, which could be targeting both CTPS and CTPS-regulating proteins, affecting the CTPS filamentation by direct ubiquitination, or indirectly, by modifying the factors which regulate the filaments. This is in agreement with our data showing that in the absence of specific ubiquitin ligases, which are in complex with CTPS, or after inhibition of ubiquitination by drugs, *S. pombe* cytoophidia are disrupted. The distinct, probably opposing roles that ubiquitination seems to play on cytoophidia formation could explain our current data in *S. pombe* DUB deletion mutants, in which cytoophidia become smaller (Figure 5).

It is known that in Drosophila ovary the presence of cytoophidia is correlated with efficient production of CTP to be used in fast synthesis of DNA and phospholipids (Strochlic et al., 2014, Wang et al., 2015). Furthermore, there are studies indicative of cooperative incorporation of the newly synthesised CTPS into the cytoophidium (Barry et al., 2014, Wang et al., 2015), suggesting that CTPS filamentation could serve as a reservoir for rapid activation, as has been previously proposed (Pai et al., 2016). To this end, our results here show that ubiquitination is an important regulator of the filamentous CTPS conformation, contributing to the dynamic nature of the structure.

Data from structural studies on human and bacterial cells revealed that CTPS cytoophidia consist of stacked tetramers of CTPS which undergo a conserved conformational cycle regulated by substrate and product binding (Lynch et al., 2017). Our study provides insights into the determinants of the process that leads to the formation of the CTPS filaments. Our results are in agreement with a model in which both possible direct effects of ubiquitination on CTPS protein itself, and indirect effects via regulation of CTPS filament regulators by ubiquitination, can determine the cytoophidium formation. We propose that the observed cytoophidia in *S. pombe* consist of bundles of self-assembled filaments which come together and form larger bundles, giving to the membraneless cytoophidia their final form that is observed in exponentially growing cells, a process that is governed by ubiquitination and deubiquitination (Figure 6).

**Fig 6.**
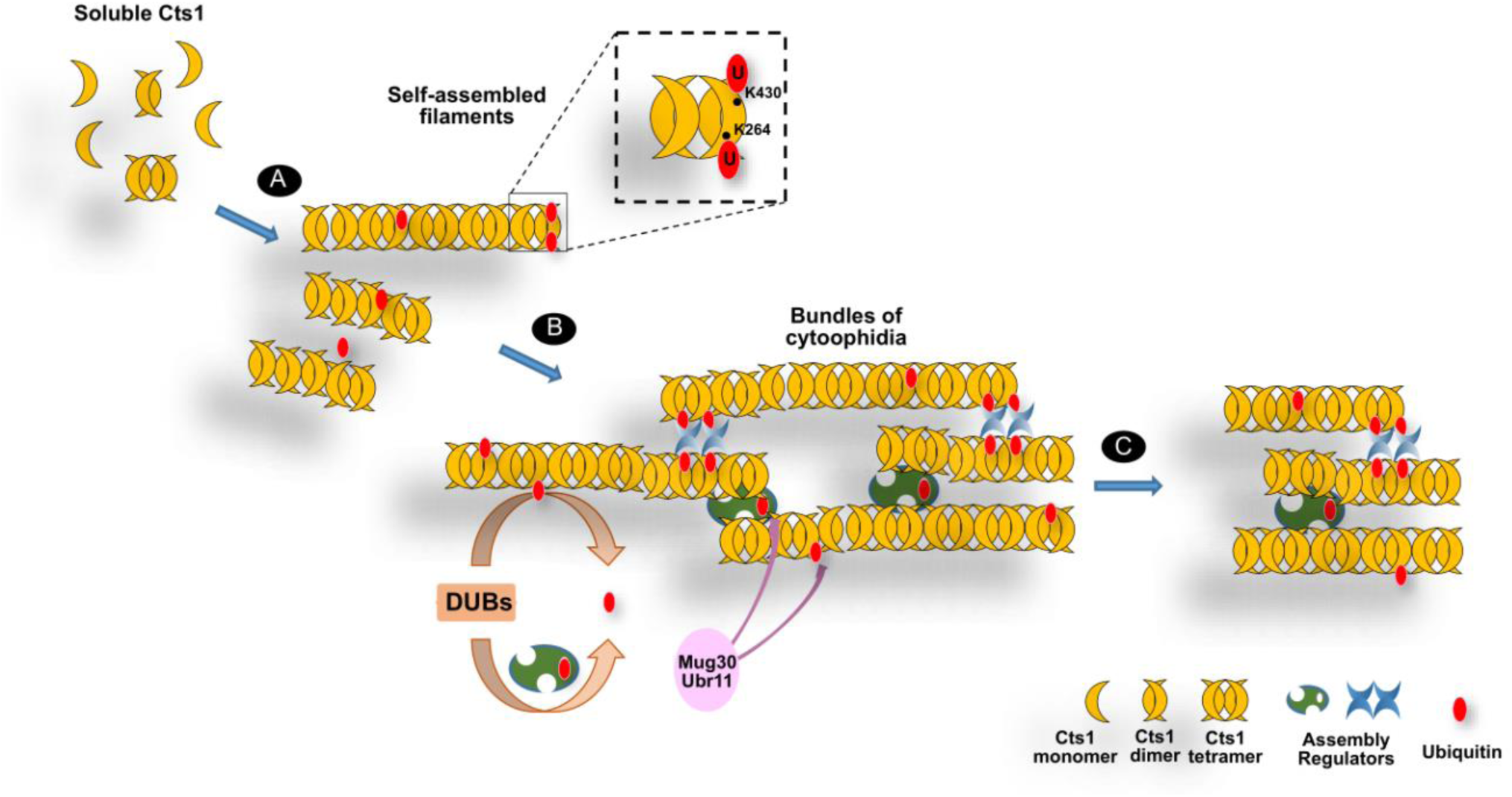
Proposed model for the role of ubiquitination in CTPS-filament assembly. Soluble CTPS protein self assembles into filaments. CTPS could be directly ubiquitinated, as our data on K430R and K264R mutants indicate, regulating its capability to form filaments (A). Ubiquitination of CTPS might promote interactions between self-assembled filaments and other possible assembly regulators, which could also be regulated by ubiquitination (B), as suggested by the changes in cytoophidia formation when the ubiquitination state of the cells is disrupted. (e.g. when ubiquitin ligation or deubiquitination do not function properly). Ubiquitin ligases, such as Mug30, and Ubr11 could affect cytoophidia formation either by directly facilitating CTPS ubiquitination or by ubiquitinating other proteins which affect filamentation indirectly. Ubiquitin hydrolases Ubp4, Ubp5, Ubp9, Ubp15 and ubiquitin-specific protease Sst2 are further regulators of CTPS compartmentation in *S. pombe.* These direct and/or indirect effects of ubiquitination on CTPS or other CTPS-regulating proteins are contributing to the observed dynamic CTPS filaments formation (C). DUB: deubiquitinating enzyme.

If CTPS filaments can be ubiquitinated, then this modification could trigger the assembly of larger bundles (Figure 6). Concurrently, it could be that regulators of CTPS filamentation, which are themselves regulated by the ubiquitination state of the cells, may contribute to the formation of dynamic CTPS filaments, as suggested by our results when ubiquitin ligation and/or deubiquitination are altered. Ubiquitin ligases such as Mug30 and Ubr11 may exhibit their regulatory effects on CTPS and/or on CTPS-regulating proteins, possibly ones in complex with CTPS. Certain deubiquitinating enzymes (Ubp4, Ubp5, Ubp9, Ubp15)) and ubiquitin-specific protease Sst2 were found to play important roles in maintaining the cytoophidia morphology in physiological conditions and contributing to their dynamic nature. Changes in the morphology or the number of cytoophidia could also be correlated with the enzymatic activity of CTPS present in the filaments; further measurements of the CTPS activity are required to test this possibility.

Given the recent research interest in studying the role of the epigenome in the Cts1 cytoophidium formation (Wang et al., 2015, Pai et al., 2016, Andreadis et al., 2019, Sun and Liu, 2019a), our study provides a basic framework for understanding the properties of CTPS as a molecule, its spatial characteristics and its function, demonstrating how a post-translational modification such as ubiquitination can define the CTPS phenotypic plasticity. It can be utilised as a reference by future studies aiming to further characterize the determinants of the regulation of CTPS filamentation by ubiquitination or other post-translational modifications (Lin et al. 2018), and would be of value in elucidating the function of CTPS cytoophidia, as well as the regulation of filaments formed by other metabolic enzymes (Noree et al., 2010, Shen et al., 2016).

## Materials and Methods

### Yeast Strains Construction and culture conditions

Cts1-YFP strain (JLL005S), was constructed as previously described (Zhang et al., 2014), after JLL003S wild type strain was transformed with linearized (SpeI) pSMUY2-Ura4-cts1-YFP plasmid. The Cts1-YFP strain has the same growth profile as the wt strain (Zhang et al., 2014). Haploid deletion strains containing Cts1-YFP were obtained by PCR-based gene targeting using pFA6A-KanMX6 or pFA6A-hphMX6 plasmids (depending on the selection marker needed) as templates for specially-designed gene-specific 80bp-long primers, as previously described (Bahler et al., 1998), followed by transformation of Cts1-YFP strains. Cts1-flag-3HA strain was constructed using the same PCR-based gene targeting method, after using pFA6A-3HA-KanMX6 plasmid as template and incorporating the flag tag in the gene-specific 80bp- long primers. The strain was verified by PCR and protein expression analyses confirming the expression of both tags. Cts1 point mutation strains were constructed by incorporating the mutated sequence leading to the K-to-R amino acid mutation into the forward primer used in the PCR-based gene targeting, using genomic DNA of a Cts1-YFP strain as template, followed by transformation of a wild type strain and colony screening. The strain bearing the point mutation was verified by sequencing. The DUB quadruple mutants were constructed after sequential mating between haploid mutants, followed by random spore analysis as previously described (Ekwall and Thon, 2017a, Ekwall and Thon, 2017b). Briefly, h+ and h- cells were streaked on YE4S agar plates and left to grow at 30°C; subsequently, fresh cells were used to set up the crosses on EMMG4S agar plates (Edinbourgh Minimal Medium containing 2% Glucose and the 4 Supplements leucine, uracil, histidine and adenine at 100μM). The plates were then left at 30°C for 2-3 days, and the formation of asci was monitored by examination under a light microscope. An inoculation loop of cells was resuspended in 1ml of 0.5% *Helix pomatia* enzyme solution and incubated overnight to break down the ascus and vegetative cells. Any remaining vegetative cells were killed by resuspending the spores in 1ml 30% ethanol. After calculating the number of spores/ml using a haemocytometer, approximately 200-1000 spores were plated on YE4S agar plates and incubated at 30°C until colonies were formed. Both the absence of each gene and its replacement by the selection cassette in the particular genetic locus were verified by PCR after genomic DNA extraction from cells derived from single colonies. Table S2 contains a list of the primers used in the construction and verification of the strains, as well as the primers used for the sequencing verifying the presence of the point mutations. Table S3 contains the strains used in this study. Standard rich media (YE4S; YES enriched with supplements leucine, uracil, histidine and adenine at 100μM) were used for the culture of *S. pombe* cells. All transformations were conducted using the lithium acetate methods (Paul Nurse lab manual). Proteasome inhibitor MG132 (474787, Merck), E1-ubiquitin ligase inhibitor PYR-41 (N2915, Sigma), and Pr619 DUB inhibitor (SML0430, Sigma) were used at the concentrations and incubation times mentioned in the text.

### Sample preparation, confocal microscopy, and statistical analysis

Cts1 cytoophidia assembly in *S. pombe* takes place in exponentially growing cells. Cells in logarithmic phase were fixed by using 16% paraformaldehyde (PFA) solution for 10 minutes while cells were still in liquid culture (final PFA concentration of 4%), washed twice with PBS buffer, and mixed with YE4S mountant (1.2% LMT agarose in YE4S, 0.1mM N-propyl gallate, to reduce phototoxicity, containing 200 ng/ml DAPI (Sigma, D9542) as DNA dye. Cells were visualized using a Zeiss LSM880 confocal microscope and photographed with Zen 2 lite (blue edition) software. ZEN 2 lite and ImageJ software were used to process the images. For the cell quantification, three independent experiments were conducted and at least 400 cells were scored manually in each of them. Two types of Cts1 cytoophidia are present in fission yeast, one longer in the cytoplasm and one shorter in the nucleus (Zhang et al., 2014). Cytoplasmic and nuclear filaments exhibited no difference; either both were present in the individual cells scored or both were absent. Three biological replicates were used to calculate the standard deviation. The percentage value of cells containing cytoophidia in each strain was calculated in each experiment, and a mean value was obtained. The statistical significance of the mean value in each mutant strain versus the mean value in the control strain was evaluated by performing a two-tailed homoscedastic t test. The cytoplasmic cytoophidia length was measured manually in at least 400 cytoophidia with the use of ImageJ software in three independent experiments. Three biological replicates were used to calculate the standard deviation. The statistical significance of the mean value of cytoophidia length for each mutant strain versus the respective mean value for the control strain was calculated by using a two-tailed homoscedastic t test.

### Western Blot Analyses

Extraction of proteins was done by using the trichloroacetic acid (TCA) method on equal number of cells for each sample, and Western blotting was performed as previously described (Ralph et al., 2006). For the detection of the YFP-tag, an anti-GFP antibody was used (Abcam ab6673, goat polyclonal). For the detection of HA tag, the HA-probe (F-7) mouse monoclonal antibody was used (Santa Cruz, sc-7392). For the detection of ubiquitin, the ubiquitin rabbit polyclonal antibody was used (10201-2-AP, Proteintech). Tubulin or actin were used as loading controls and the detection was done by using anti-tubulin antibody (Sigma T5168, mouse monoclonal) or anti-actin antibody (ab14128, Abcam). The secondary antibodies used were Peroxidase-AffiniPure Donkey anti-goat IgG (Jackson ImmunoResearch, 705-035-147), anti-mouse IgG HRP-linked (Cell Signalling Technology, 7076), and peroxidase anti-rabbit IgG (111-035-144, AffiniPure). The protein signal was detected by using SuperSignal West Pico Chemiluminescence Substrate (Thermo Scientific, 34087), and exposing the membrane in an Amersham Imager 600 detector. Quantification of the protein bands was done with the use of ImageJ software.

### Co-immunoprecipitation assay

To detect *in vivo* interactions between CTPS and ubiquitin in wild type and in mutant cells we conducted co-immunoprecipitation assays. Cts1-flag-3HA or Cts1-YFP cells were cultured in rich YE4S medium until they reached exponential phase (OD600=0.4) and 5×10^7^ cells were collected. The cells were then suspended in Buffer A (50mM Hepes-KOH pH=7.5; 5mM MgAc; 100mM KAc; 0.1% NP40; DTT 1mM, BSA 0.5mg/ml), and protease inhibitor mixture (Sigma, P8215) was added, along with glass beads. The cells were then beaten with the glass beads for 5 min at 4°C on a Vortex machine (30 seconds of beating followed by 30 seconds of resting time). When the glass beads precipitated due to gravity, the supernatant was collected. The extract was then centrifuged at 4°C for 15mins at 13000rpm and the supernatant was transferred to a fresh tube. Approximately 1/10 of the volume was kept as Input sample, while the rest of the extract was immunoprecipitated using 2μg α-ubiquitin rabbit polyclonal antibody (10201-2-AP, Proteintech) after overnight incubation at 4°C. The extract was then incubated with A/G magnetic beads (GeneScript, L00277), following the manufacturer’s protocol. The elution was performed by suspending the magnetic beads in Laemmli buffer and boiling the mixture for 5 minutes, before performing SDS-PAGE electrophoresis of the supernatant, as described in the Western blot analyses section. For the detection of the YFP-tag, an anti-GFP antibody was used (Abcam ab6673, goat polyclonal). For the detection of HA tag the HA-probe (F-7) mouse monoclonal antibody was used (Santa Cruz, sc-7392). For the detection of ubiquitin, the ubiquitin rabbit polyclonal antibody was used (10201-2-AP, Proteintech). The secondary antibodies used were Peroxidase-AffiniPure Donkey anti-goat IgG (Jackson ImmunoResearch, 705-035-147), anti-mouse IgG HRP-linked (Cell Signalling Technology, 7076), and peroxidase anti-rabbit IgG (111-035-144, AffiniPure). The experiments were performed in triplicate. For the case of Cts1-YFP the intensity of the bands was quantified by using ImageJ software.

### Tandem affinity purification (TAP) and Liquid chromatography - Mass spectrometry (LC-MS) Analyses

We used a Cts1-flag-3HA strain for our TAP and LC-MS analyses. A wild type strain was used as reference. For the TAP assay we used the FLAG-HA tandem affinity purification kit manufactured by Sigma (TP0010) and performed the protocol according to the manufacturer’s specifications. Briefly, Cts1-flag-3HA cells were cultured in rich media (YE4S) until they reached exponential phase. 50×10^7^ cells were collected and the cell lysate was extracted, as described in the co-IP protocol. The cell lysate was then incubated with anti-FLAG M2 resin rotating overnight at 4°C, ensuring efficient binding. The supernatant was then removed carefully, and the resin was washed with RIPA buffer (Sigma, R0278) containing protease inhibitors (Sigma, P8215), in order to remove any unbound protein. The first elution of the protein complex bound on the resin was done by using 3XFLAG peptide, and in a following step the eluate was bound to anti-HA resin slurry. In the second elution of the protein complex, the anti-HA slurry was washed with TBS (50mM Tris-Cl, ph=7.5; 150mM NaCl), to remove any unbound protein. The final elution was done by using TBS with 100mM ammonium bicarbonate, and the sample was subsequently digested with trypsin overnight.

Peptides were then separated and analyzed on an Easy-nLC 1000 system coupled to a Q Exactive HF (both - Thermo Scientific). About 2 µg of peptides were separated in an home-made column (75 µm x 15 cm) packed with C18 AQ (5 µm, 300Å, Michrom BioResources, Auburn, CA, USA) at a flow rate of 300 nL/min. Mobile phase A (0.1% formic acid in 2% ACN) and mobile phase B (0.1% formic acid in 98% ACN) were used to establish a 60 min gradient comprised of 2 min of 5% B, 40 min of 5-30% B, 6 min of 30-45% B, 2 min of 45-90% B and 10 min of 90% B. Peptides were then ionized by electrospray at 2.3 kV. A full MS spectrum (300-1800 m/z range) was acquired at a resolution of 120,000 at m/z 200 and a maximum ion accumulation time of 20 ms. Dynamic exclusion was set to 30 s. Resolution for HCD MS/MS spectra was set to 30,000 at m/z 200. The AGC setting of MS and MS2 were set at 3E6 and 1E5, respectively. The 20 most intense ions above a 1.3E4 counts threshold were selected for fragmentation by HCD with a maximum ion accumulation time of 80 ms. Isolation width of 1.6 m/z units was used for MS2. Single and unassigned charged ions were excluded from MS/MS. For HCD, normalized collision energy was set to 30%. The underfill ratio was defined as 1%.

The raw data were processed and searched with MaxQuant 1.5.4.1 with MS tolerance of 4.5 ppm, and MS/MS tolerance of 20 ppm. The Uniprot *Schizosaccharomyces pombe* proteome database and the database for proteomics contaminants from MaxQuant were used for database searches. Reversed database searches were used to evaluate false discovery rate (FDR) of peptide and protein identifications. Two missed cleavage sites of trypsin were allowed. Oxidation (M), Acetyl (Protein N-term), deamidation (NQ) and GGE (K) were set as variable modifications. The FDR of both peptide identification and protein identification is set to be 1% (Elias and Gygi, 2007). The option of “Second peptides”, “Match between runs” and “Dependent peptides” was enabled. Label-free quantification was used to quantify the difference of protein abundances between different samples (Cox and Mann, 2008, Cox et al., 2011).

### 3D modelling of CTPS and ubiquitination target prediction

Based on the resolved CTP synthase three-dimensional conformation, the mammalian CTP synthase amino acid sequence was used as a template in Phyre2 software (Kelley et al., 2015) to obtain the PDB formatted model of the *S. pombe* CTP synthase, which was subsequently studied with EzMol software (Reynolds et al., 2018), in order to model the three-dimensional conformation of *S. pombe* CTPS and predict the space occupied by the residues of interest. For the prediction of potential ubiquitination-targeted amino acids on CTPS we used the fission yeast CTPS protein sequence as a template in Ubipred software (Radivojac et al., 2010).

### Gene Ontology (GO) Analyses

For the Gene Ontology analyses, the Cytoscape software platform (Cline et al., 2007) was utilised to visualise the potential networks in which the proteins in complex with CTPS participate. In order to determine the gene ontology (GO) terms that are significantly overrepresented, the biological networks gene ontology (BiNGO) tool (Maere et al., 2005) was used.

## Acknowledgements

The authors would like to thank Han Ying for technical assistance in the LC-MC facility at ShanghaiTech University.

## Data Availability

The MS raw data are deposited on Figshare public database (https://figshare.com/, DOI: 10.6084/m9.figshare.14071499), containing details on the number of peptides assigned for each protein and on the sequence coverage for each hit.

## Supplementary Information

**Table S1.** Proteins in complex with S. pombe CTPS. A Cts1-FLAG-HA double tagged protein was constructed, and the cells were grown in rich YE4S medium until they reached exponential phase (OD600=0.8). After the preparation of the cell lysate, tandem affinity purification by two sequential elution steps followed. Initially, the cell lysate was bound to anti-FLAG resin and competitive elution with FLAG peptide was conducted. Then, the solution was bound to anti-HA resin, before a second elution ensured that the Cts1 protein complex was efficiently purified. The purified protein complex was analysed by liquid chromatography-mass spectrometry (LC-MS) and the detected proteins present in at least two biological replicates are presented in the table, after comparison with a reference sample. The proteins found to be in complex with CTPS are presented in the table based on the major biological processes in which they participate; structural (in yellow), metabolism (in blue), transport (in green), and stress-related (in pink). The experiment was performed at least in triplicate.

| Protein | Description |
| --- | --- |
| Act1 | actin |
| Psm3 | Structural maintenance of chromosome protein 3 |
| Fim1 | fimbrin |
| Mbo1/Mto 1 | Microtubule organizer protein 1 |
| Nda2/Tub 1 | nda2 / tub1 tubulin alpha-2 chain;Tubulin alpha-1 chain |
| Cyr1 | Adenylate cyclase |
| Lid2 | Lid2 complex component lid2 |
| Mas2 | Probable mitochondrial-processing peptidase subunit alpha |
| SPCC126.12 | GTP cyclohydrolase (predicted), Protein NIF3 homolog |
| Ufe1 | syntaxin ufe1 |
| Cao1 | Copper amine oxidase 1 |
| Coq11 | NADH dehydrogenase [ubiquinone] complex I, assembly factor 7 homolog |
| Ubp5 | Probable ubiquitin carboxyl-terminal hydrolase 5 |
| Ade8 | Adenylosuccinate lyase |
| Tif225 | translation initiation factor eIF2B epsilon subunit |
| Npg1 | GTPase regulator Rng2-like (predicted) |
| Mug30 | HECT-type ubiquitin-protein ligase E3 |
| Ubr11 | ubiquitin-protein ligase E3 (UBR family) Ubr11 |
| Aly1 | SPBC839.02 Putative arrestin-related trafficking adapter |
| Ski7 | SPAP8A3.05 Ski complex interacting GTPase Ski7 |
| Kap109 | Importin-alpha re-exporter |
| Kap114 | Importin subunit beta-5 |
| Can1 | plasma membrane arginine/lysine transmembrane transporter Can1 |
| Swo1/Hsp90 | Heat shock Protein 90 homolog |

**Table S2.**
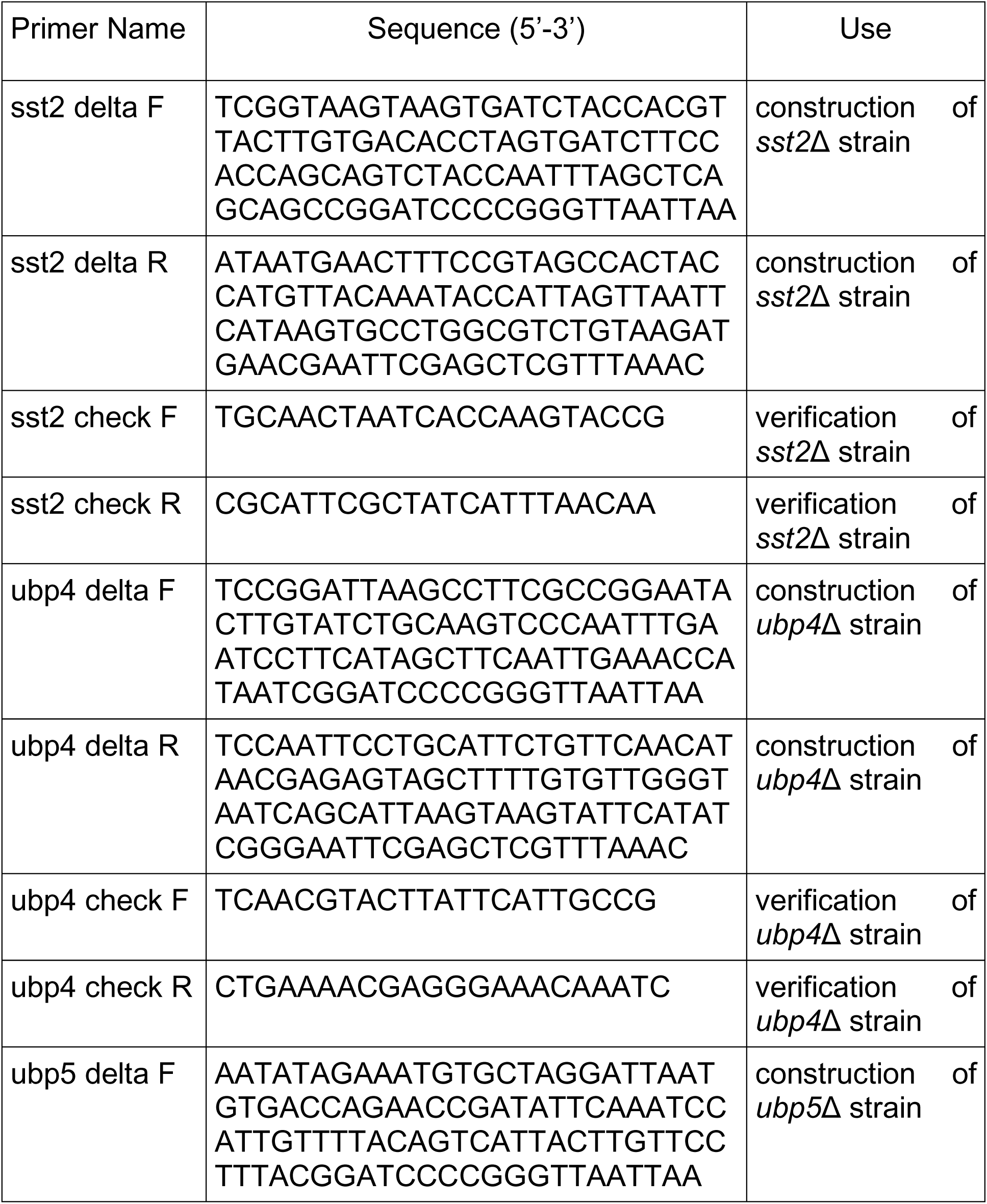
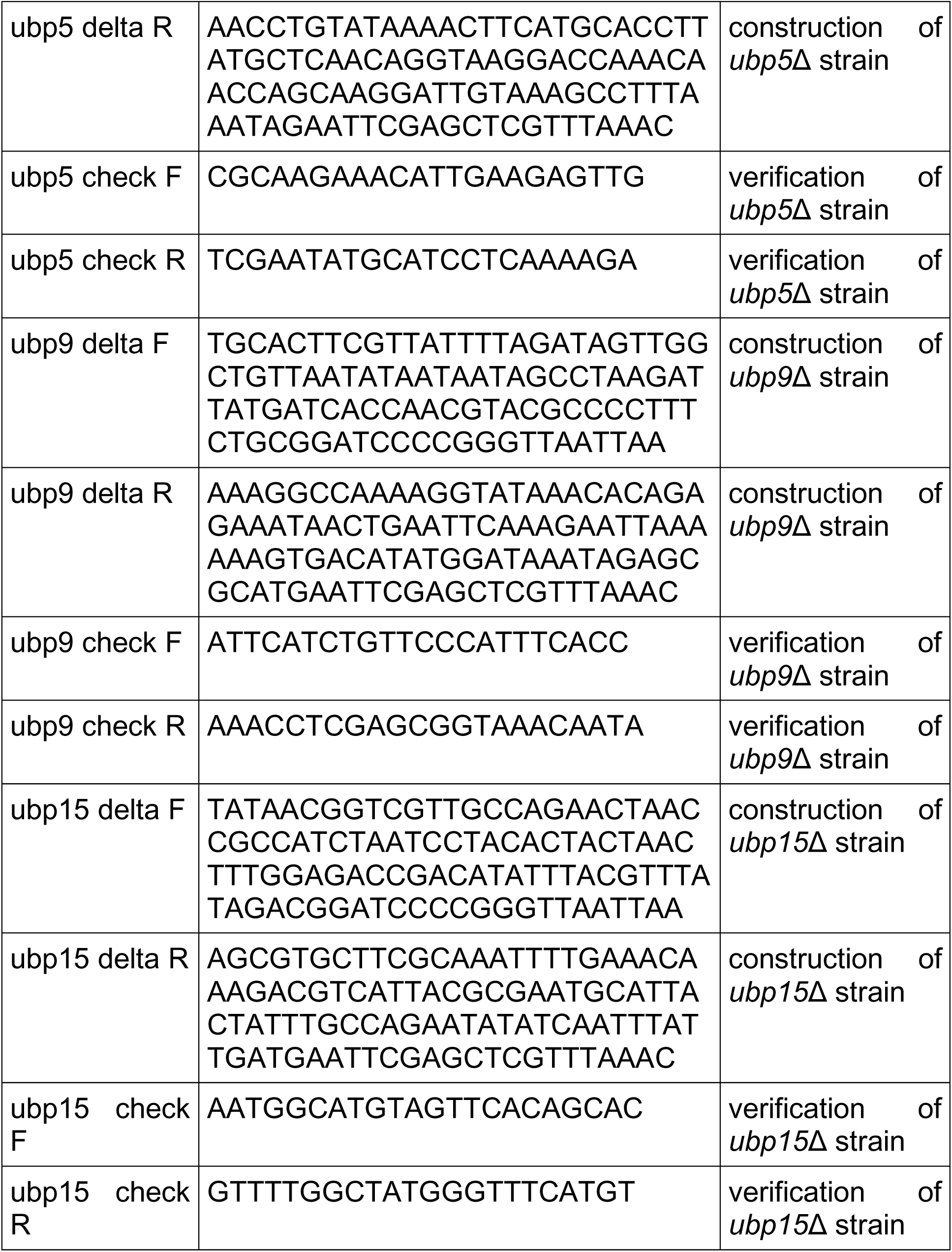
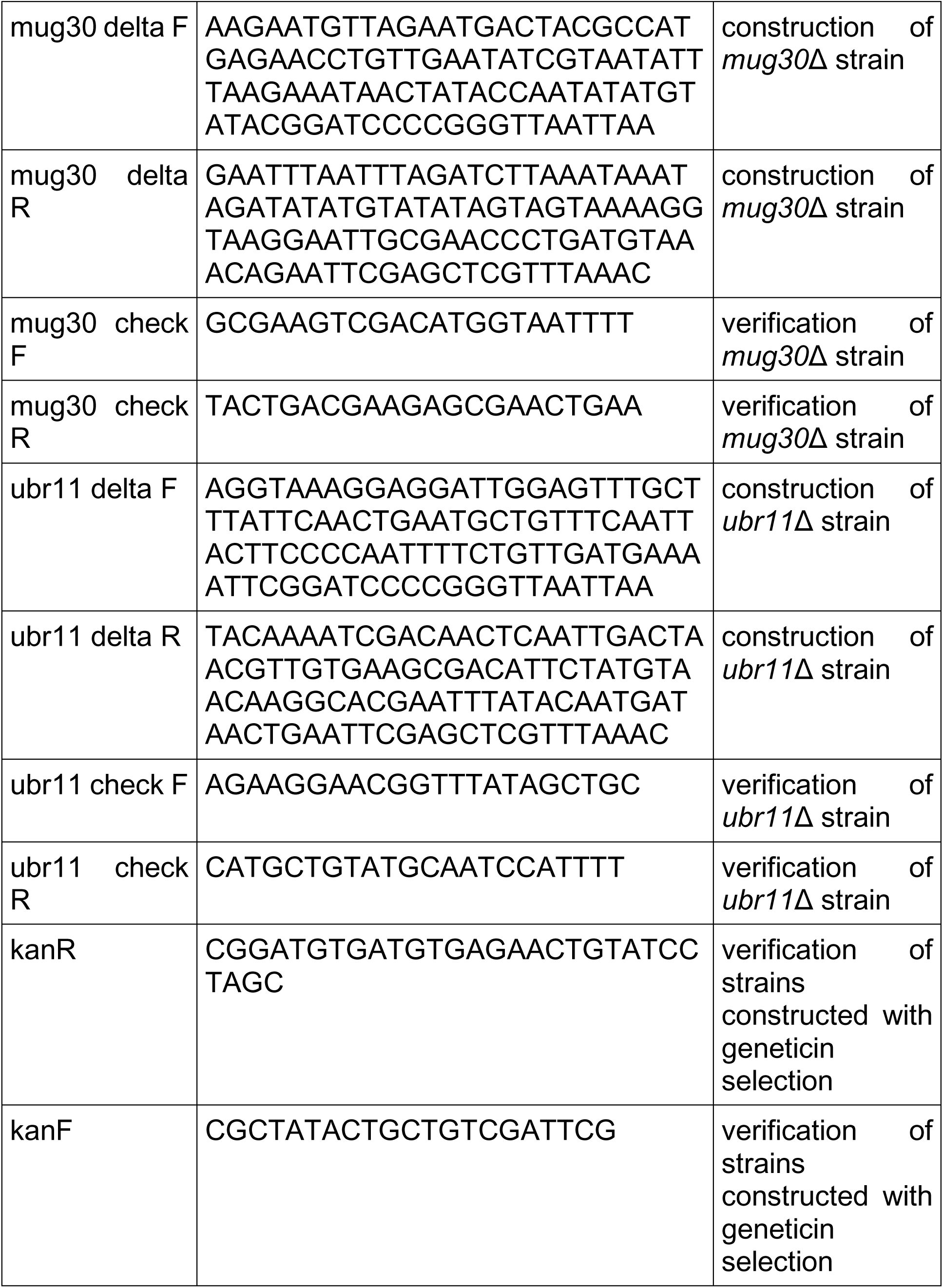
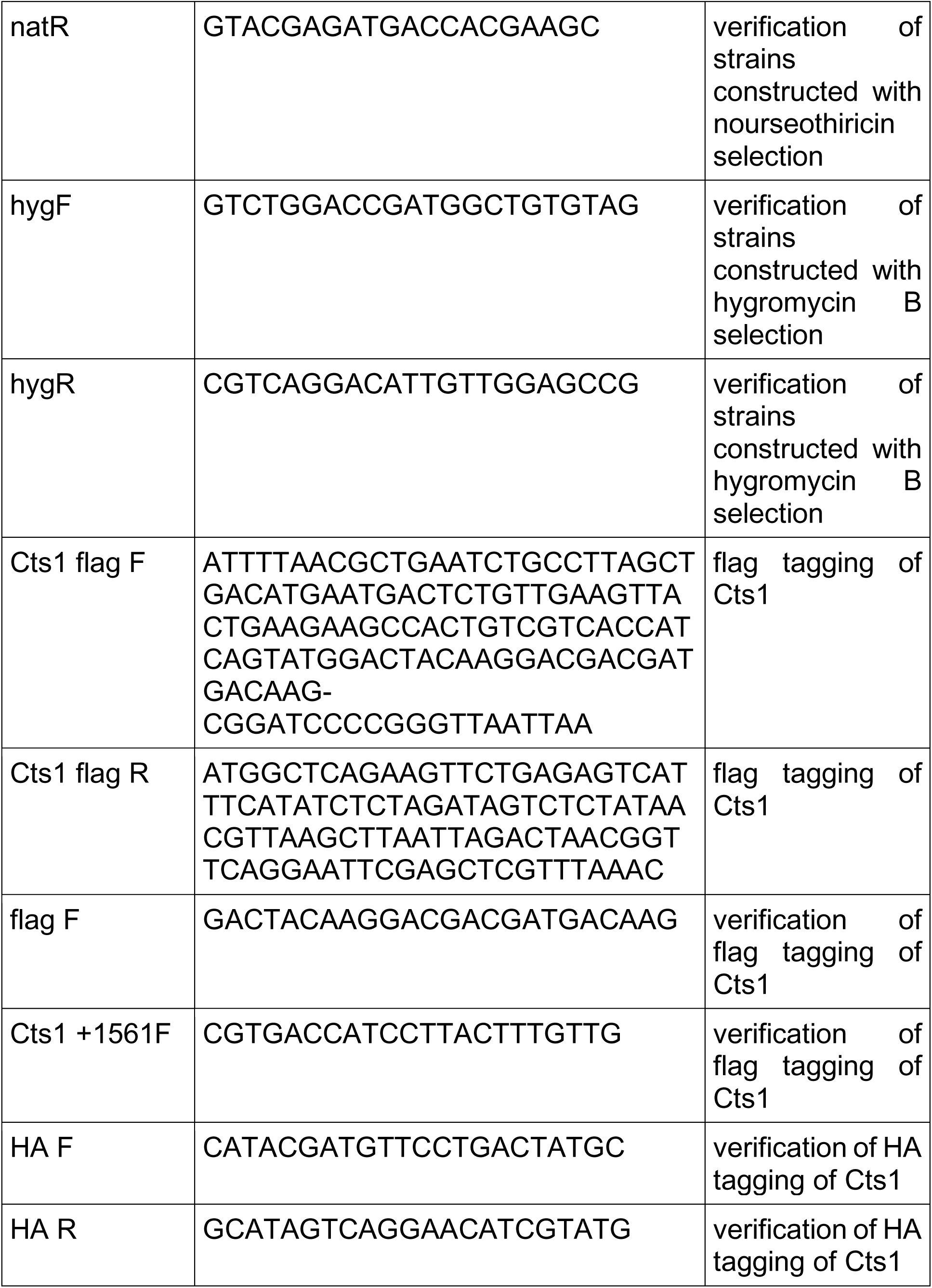
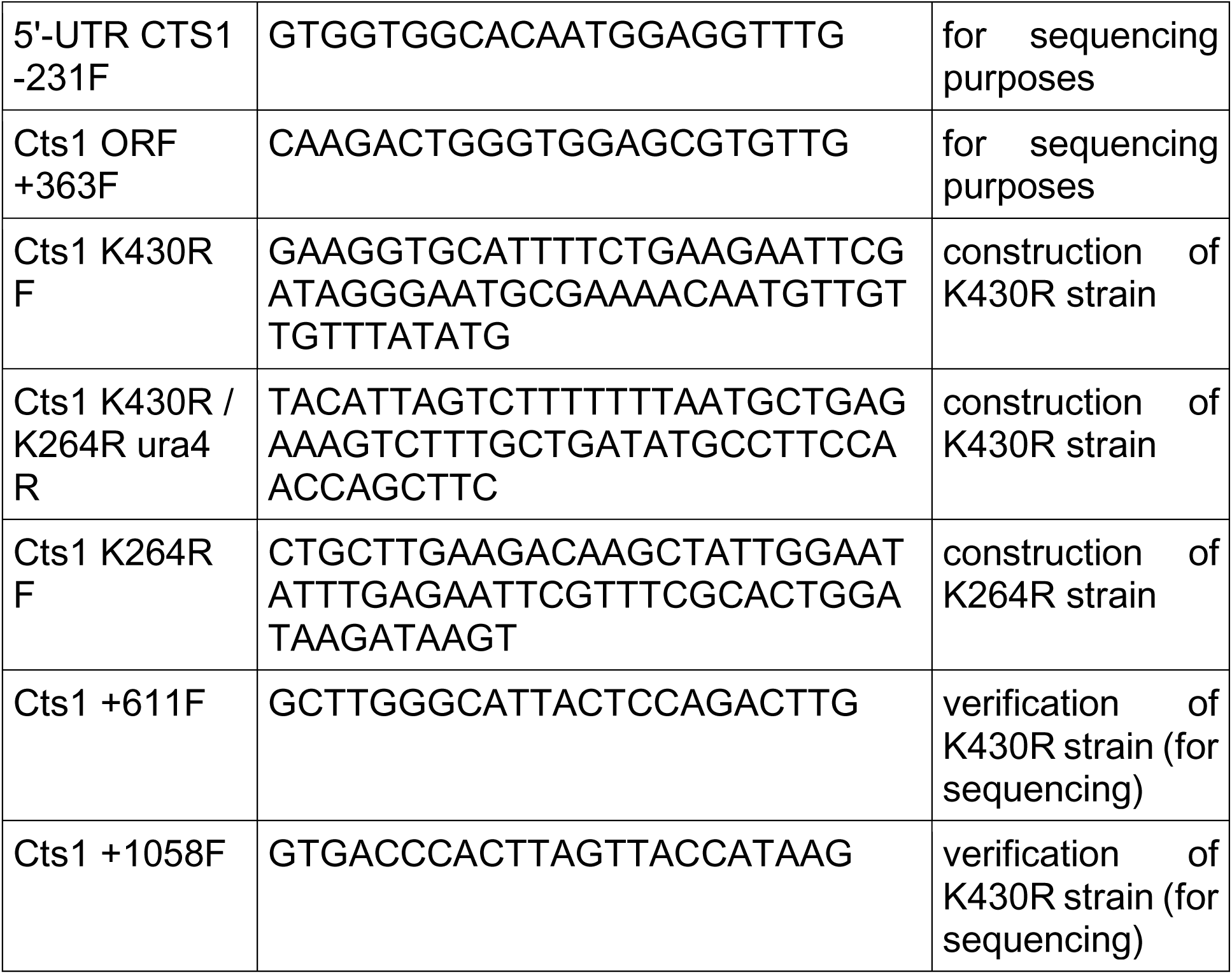
Primers used in this study.

**Table S3.** S. pombe strains used in this study.

| Strain Number | Genotype |
| --- | --- |
| JLL003S | h- *his3*-D1 *ura4*-D18 *leu1*-32 *ade6*-M216 |
| JLL005S | h+ *cts1*-YFP (*ura4*+) *ura4*-D18 *leu1*-32 *ade6*-M216 |
| JLL53S | h+ *cts1*-YFP (*ura4*+) *sst2*::KanMX6 *ura4*-D18 *leu1*-32 *ade6*-M216 |
| JLL63S | h+ *cts1*-YFP (*ura4*+) *ubp4*::KanMX6 *ura4*-D18 *leu1*-32 *ade6*-M216 |
| JLL63S | h+ *cts1*-YFP (*ura4*+) *ubp5*::KanMX6 *ura4*-D18 *leu1*-32 *ade6*-M216 |
| JLL59S | h- *cts1*-YFP (*ura4*+) *ubp9*::KanMX6 *ura4*-D18 *leu1*-32 *ade6*-M216 |
| JLL60S | h- *cts1*-YFP (*ura4*+) *ubp15*::KanMX6 *ura4*-D18 *leu1*-32 *ade6*-M216 |
| JLL87S | h- *cts1*-YFP (*ura4*+) *sst2*::KanMX6 *ubp4*::KanMX6 *ubp9*::KanMX6 *ubp15*::KanMX6 *ura4*-D18 *leu1*-32 *ade6*-M216 |
| JLL88S | h+ *cts1*-YFP (*ura4*+) *sst2*::KanMX6 *ubp4*::KanMX6 *ubp5*::KanMX6 *ubp9*::KanMX6 *ura4*-D18 *leu1*-32 *ade6*-M216 |
| JLL105S | h+ *cts1*-flag-3HA (KanMX6) *ura4*-D18 *leu1*-32 *ade6*-M216 |
| JLL114S | h- *cts1*-YFP (*ura4*+) *mug30*::KanMX6 *ura4*-D18 *leu1*-32 *ade6*-M216 |
| JLL115S | h- *cts1*-YFP (*ura4*+) *ubr11*::hyg *ura4*-D18 *leu1*-32 *ade6*-M216 |
| JLL174S | h+ *cts1*-YFP (*ura4*+) *cts1* K430R *ura4*-D18 *leu1*-32 *ade6*-M216 |
| JLL175S | h+ *cts1*-YFP (*ura4*+) *cts1* K264R *ura4*-D18 *leu1*-32 *ade6*-M216 |

**Fig S1.**
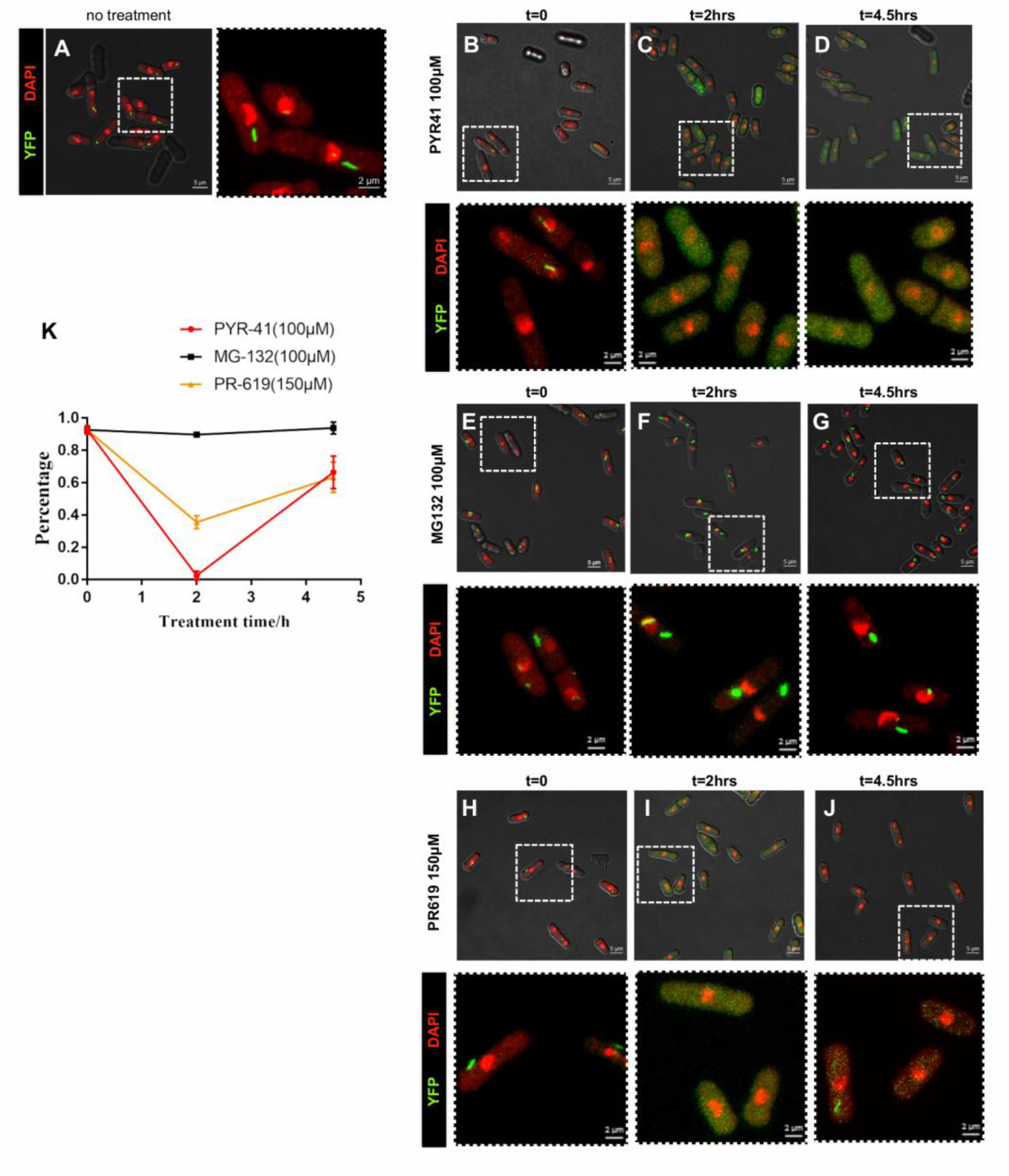
Cytoophidia in *S. pombe* recover after prolonged drug treatment. (A) CTP synthase fused to YFP was expressed endogenously under the control of *Cts1* native promoter. *S. pombe* control cells were cultured until exponential phase (OD600=0.4) in YE4S rich medium with the addition of 0.4% DMSO for 2h, fixed, and observed by confocal microscopy. An overlap image of brightfield (white), YFP (green), and DAPI (red) is presented on the left part (scale bar=5μm), while a magnified image of the dotted rectangle is presented on the right with an overlap of the YFP and DAPI channels for clarity (scale bar=2μm). (B-D) Exponentially growing *S. pombe* cells (OD600=0.4) were treated with 100μM PYR-41 for 2h and 4.5h, fixed, and observed by confocal microscopy (t=0: the time point at which the drug was added). (E-G) Exponentially growing *S. pombe* cells (OD600=0.4) were treated with 100μM MG-132 for 2h and 4.5h, fixed, and observed by confocal microscopy (t=0: the time point at which the drug was added). (H-J) Exponentially growing *S. pombe* cells (OD600=0.4) were treated with 150μM PR-619 for 2h, fixed, and observed by confocal microscopy (t=0: the time point at which the drug was added). (K) Plot demonstrating the changes in the percentage of cells containing cytoophidia, after treatment with PYR-41, MG-132 and PR-619 for 2 or 4.5h. Error bars show the mean ± S.D.: as calculated from three independent experiments (>400 cells were manually counted per strain per trial).

**Fig S2.**
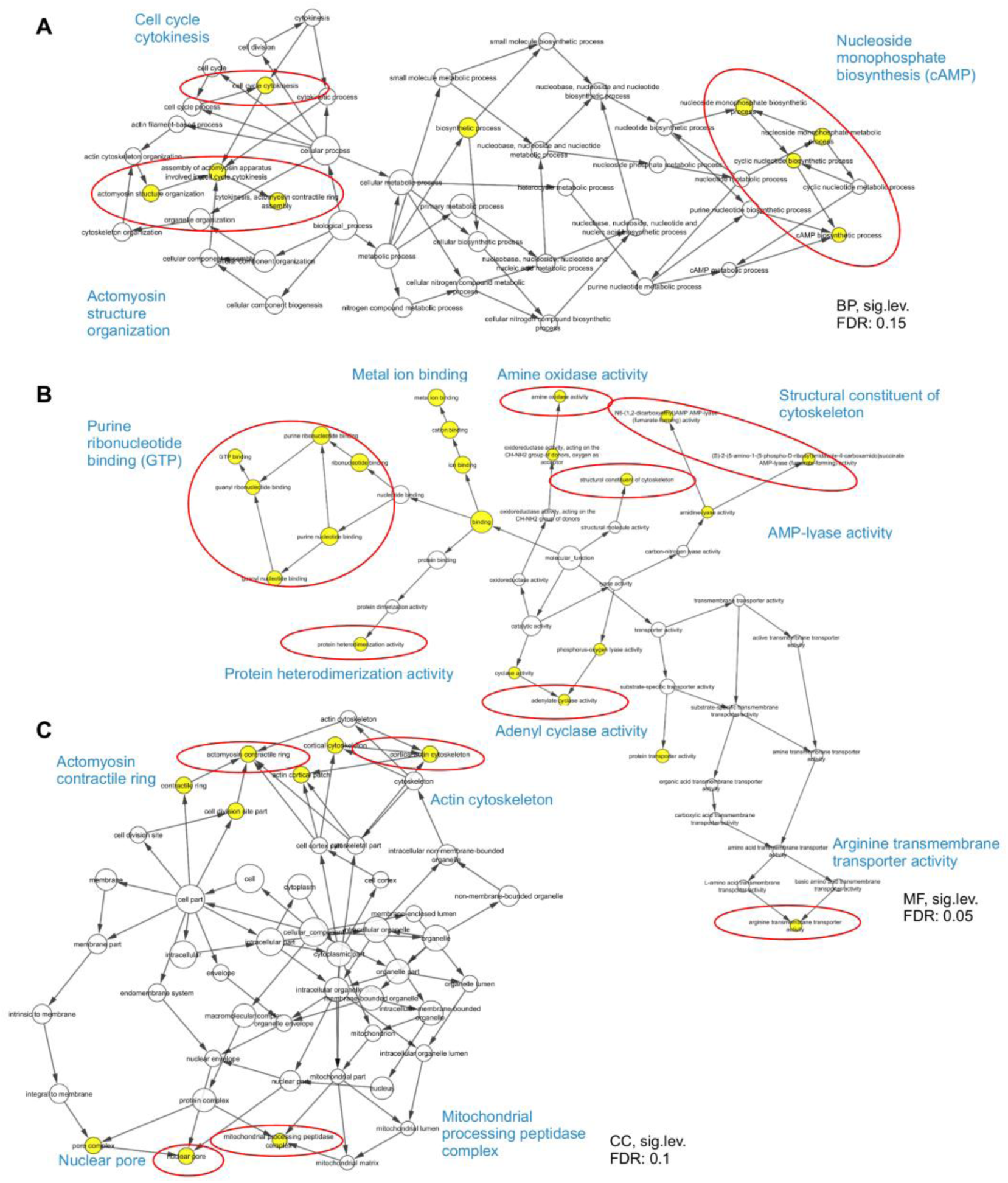
Gene ontology analysis of *S. pombe* proteins in complex with CTPS. Visualisation of gene ontology (GO) terms that are significantly overrepresented in the proteins found in complex with CTPS in *S. pombe.* Networks of GO terms related to the biological process (A), molecular function (B), and cellular component (C), in which the group of proteins are participating are shown. Yellow colour in the circles indicates higher statistical significance. Highlighted by red circles are significant characteristic GO terms and a simplified description of the overrepresented groups is noted in blue font. BP: Biological Process, MF: Molecular Function, CC: Cellular Component, sig. lev.: significance level, FDR: False Discovery Rate.

**Fig S3.**
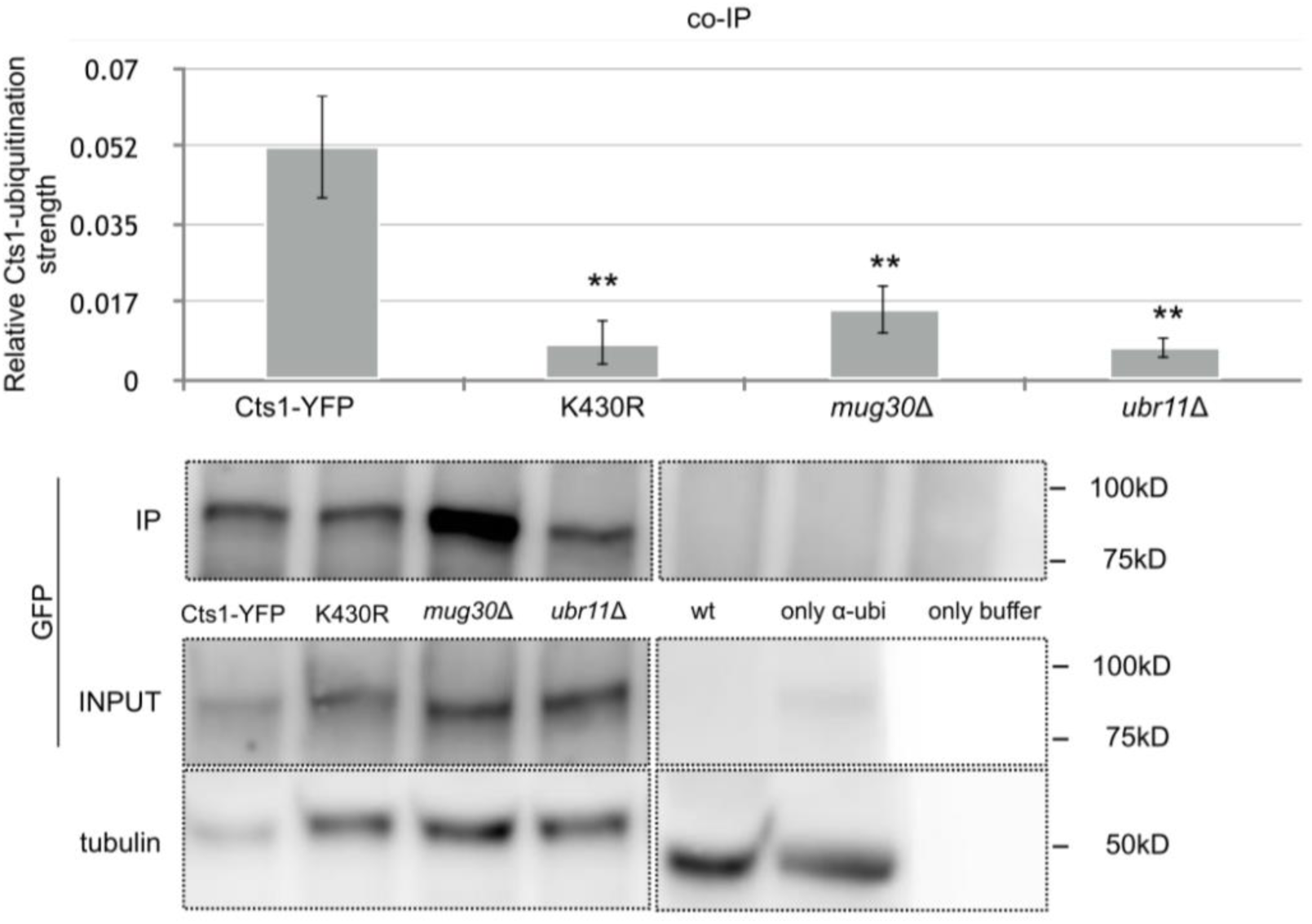
Co-immunoprecipitation of CTPS with ubiquitin is attenuated in ubiquitin ligases mutants. Cts1-YFP cells, as well as ubiquitin ligase deletion mutants and K430R strains, all in a Cts1-YFP background, were cultured in rich YE4S medium until they reached exponential phase (OD_600_=0.4). The protein extract was subsequently immunoprecipitated using α-ubiquitin antibody overnight, followed by incubation with A/G beads. The samples were then electrophoresed on SDS-PAGE and the membrane probed by α-GFP antibody. The calculated average Cts1-YFP signal obtained from two independent immunopreciptations is shown on the graph. A representative image for the Western blot analysis is shown below for the immunoprecipitated samples (IPs) and input samples (INPUTs). Control samples included one for the α-ubiquitin antibody background (α-ubi ctrl), and one for the A/G beads background (A/G beads ctrl), as well as the wt strain (no tag). The quantification of the Cts1 IP signals was normalised over the relevant Input signal. Error bars show the mean ± S.D.: as calculated from two independent experiments (**: p<0.01, n.s.: not significant). Dotted lines indicate the areas of the membrane that have been cut out.

**Fig S4.**
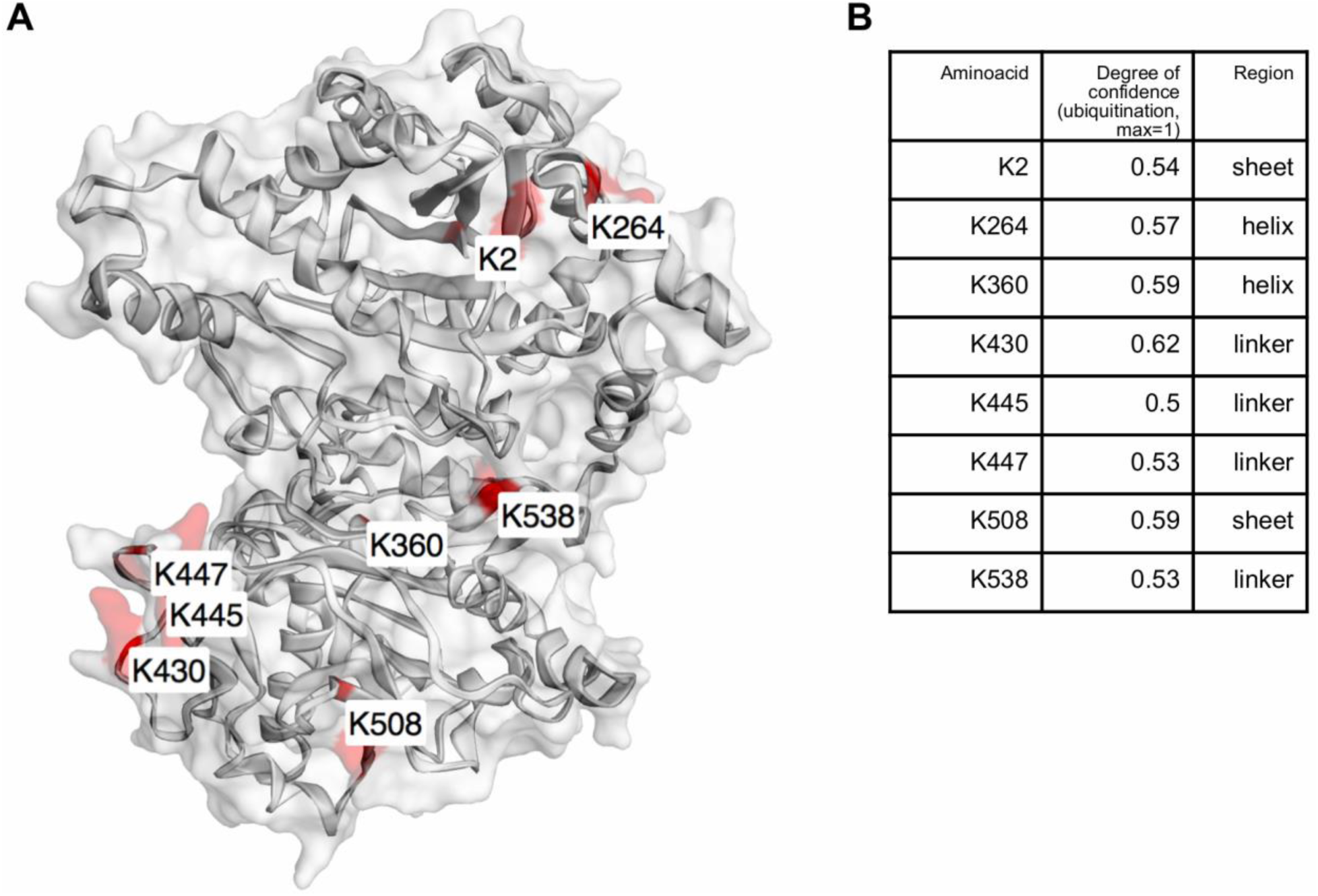
Potential ubiquitination targets on *S.pombe* CTPS. (Α) Three-dimensional conformation of *S. pombe* using Phyre2 and EzMol software, based on the solved conformation of mammalian CTPS. *In silico* analysis utilizing UbiPred software tested the probability of CTPS lysine residues to be ubiquitinated; the positioning of the eight amino acids with the highest degree of confidence is indicated. (B) Table showing the degree of confidence for each of the potential lysine ubiquitination targets on CTPS, along with their domain positioning on the three-dimensional model of *S. pombe* CTPS.

